# Systematic development of degradable polyester biomaterials via ring-opening copolymerization of succinic anhydride and epoxides

**DOI:** 10.1101/2025.10.21.683741

**Authors:** Sara C. Murrin, Kaitlyn E. Woodworth, Brenden Wheeler, Zachary S.C.S. Froom, Alison J. Scott, Locke Davenport Huyer

## Abstract

Degradable polyester materials are widely utilized in medicine as resorbable sutures, implantable devices, and drug delivery. These applications require precise and tunable degradation control; predictable number-average molecular weight (*M̅_n_*), narrow polydispersity (PDI), and diverse material properties define polyester utility, which are not easily achieved through well-established synthesis approaches. Ring-opening copolymerization (ROCOP) provides reproducible *M̅_n_* control, narrow PDI, and expands monomer diversity. In this work, poly(cyclohexene succinate) (PCS) and poly(propylene succinate) (PPS) were synthesized through a central composite design of experiments approach, systematically varying anhydride:epoxide ratio, monomer:catalyst ratio, reaction temperature, and reaction time. Reduced synthesis factor-response models explained significant variation for all characterized properties relevant to degradation control. PCS and PPS readily degraded under base-catalyzed hydrolysis conditions with significantly higher mass loss in PPS materials compared to PCS, highlighting the monomer selection influence in degradation behaviour. These findings highlight the potential for ROCOP to generate degradable biomaterials with reproducible material properties in application-specific biomedical use.

## 1 INTRODUCTION

Polyester materials have seen considerable application in medicine (e.g. resorbable sutures^1^, implantable devices^2^, and controlled drug delivery^3^) owing to their tunable material properties and capacity for *in situ* resorption, which eliminates the need for secondary removal. The rate of *in situ* degradation, which occurs primarily through hydrolysis of ester linkages, is strongly influenced by polymer composition and structure^4^. At the molecular level, monomer selection determines electrophilic properties of the ester linkage, polymer hydrophilicity, and underlies chain interactions that define the general resorption timeline (i.e., months vs. years)^5^. Subsequently, differences in macromolecular features, including number-average molecular weight (*M̅_n_*), polydispersity (PDI), and glass transition temperature (T_g_), which dictate chain length and entanglement, further define degradable material lifespan and performance. Rational design of degradable polyesters can be achieved through deliberate manipulation of these characteristics for predictable and application-specific performance.

Synthesis of polyesters relevant to degradable biomaterial applications has primarily been achieved through polycondensation or ring-opening polymerization approaches^6–8^. Step-growth polycondensation of acid and alcohol functionalities yields polyester materials of diverse composition achieved through homo- or co-polymerization of α-hydroxyacids (e.g., lactic acid, glycolic acid), as well as diverse formulations through copolymerization of multi-functional carboxylic acids with comparable alcohols^4^. While effective at yielding a broad range of polyester material properties, this approach is limited by poor control over key material properties (i.e., *M̅_n_* and PDI), which can further vary between synthesis batches^9,10^. As a result of these challenges, clinically approved polyester technologies have largely relied on chain-growth ring opening polymerization of cyclic lactones^11^ (i.e., lactide, glycolide, ε-caprolactone), which achieves a comparably predictable *M̅_n_* and narrow PDI controlled by synthesis factors (e.g., monomer:catalyst ratio, initiator concentration)^12^. Improved property precision has enabled predictable degradation control, supporting the use of α-hydroxyacid-based polyesters in applications such as resorbable sutures and drug delivery systems that require defined degradation timelines^13^.

Recent advances in catalysis and monomer combinations have expanded ring opening polymerization to achieve alternating copolymerization of cyclic anhydrides and epoxides (ring opening copolymerization; ROCOP). This strategy incorporates carboxylic acid derivatives (i.e., succinic anhydride (SA)^14–19^, itaconic anhydride (IA)^14^, and citraconic anhydride (CA)^20^) that were traditionally limited to condensation approaches^21–23^ with various epoxides, such as propylene oxide (PO), butylene oxide, and cyclohexene oxide (CHO)^24^. As with traditional lactone-based ring opening polymerization, ROCOP-derived material properties can be manipulated through synthesis input parameters including monomer feed ratios^25,26^, catalyst concentration^19,27,28^, and reaction conditions^29,30^. Together, these advances provide a new avenue in polyester biomaterial development that couples the compositional diversity of copolymerization with the precision of chain-growth ring opening polymerization.

In this work, we investigate the ROCOP synthesis of SA with PO and CHO to synthesize poly(cyclohexene succinate) (PCS) and poly(propylene succinate) (PPS). To appreciate the relationships between polyester material properties and synthesis parameters, we employed a statistically informed central composite design of experiments (CCD) approach^31^, systematically varying anhydride:epoxide (A:E) ratio, monomer:catalyst (M:C) ratio, reaction temperature, and reaction time. Reduced synthesis factor-response models explained significant variation for all characterized properties relevant to degradation control. PCS and PPS readily degraded under base-catalyzed and neutral hydrolysis conditions, with significantly higher mass loss in PPS materials compared to PCS, highlighting the monomer selection influence in degradation behaviour. ROCOP may provide a novel strategy to generate degradable biomaterials with reproducible material properties in application-specific biomedical use.

## 2 MATERIALS AND METHODS

### 2.1 Materials

SA (95.0% purity), CHO (98.0% purity), PO (99.0% purity), and bis(triphenylphosphine)iminium chloride (PPNCl) (98.0% purity) were purchased from TCI Chemicals (Portland, United States) and stored under nitrogen. Calcium hydride (95.0% purity, stored under nitrogen), Cr(III) salen (stored under nitrogen), HPLC-grade methanol (MeOH) (99.8% purity), HPLC-grade dichloromethane (DCM) (99.8% purity), deuterated chloroform (CdCl_3_, 99.8% purity), butylhydroxytoluene (BHT, 99.0% purity), dimethyl sulfoxide (DMSO), phosphate buffer (1M) and 4Å molecular sieves were purchased from Sigma Aldrich (St. Louis, United States). Toluene (99.5% purity) was purchased from VWR Chemicals (Radnor, United States). Isopropyl alcohol (IPA, 99.5% purity), hexanes (98.5% purity), sodium hydroxide (NaOH, 97% purity), and HPLC-grade tetrahydrofuran (THF, 99.9% purity, stabilized with 250ppm BHT) were purchased from Fisher Scientific (Waltham, United States). Anhydrous ethanol (100% purity) was sourced from Commercial Alcohols, Greenfield Global (Mississauga, Canada). Narrow and EasiVial polystyrene standards were purchased from Agilent Technologies (Santa Clara, United States). Thermal analysis polystyrene standard was purchased from Polymer Source Inc. (Montreal, Canada). Thermal analysis indium standard was purchased from TA Instruments (New Castle, DE, United States). Dulbecco’s modified eagle medium with L-glutamine (DMEM), heat inactivated fetal bovine serum (FBS), Dulbecco’s phosphate-buffered saline (DPBS), and penicillin/streptomycin solution were purchased from Corning (Corning, NY). NIH3T3 cells were purchased from ATCC (Manassas, VA). Hoechst 33342, 5-(and-6)-carboxyfluorescein diacetate, succinimidyl ester (CFDA-SE), and propidium iodide (PI) were purchased from ThermoFisher Scientific (Waltham, United States). All reagents were used as received unless otherwise indicated.

### 2.2 Polyester Synthesis and Extraction

Polymer synthesis was performed with monomer reagent quantities, polymerization time, and polymerization temperature specified according to a CCD (**Table 1**). SA and epoxide (PO, CHO) monomers were purified via sublimation or distillation, respectively, prior to use under inert atmosphere conditions. SA, Cr(III) salen catalyst, PPNCl, and epoxide were combined with toluene (4.0 mL) in a pressure-safe 20mL vial under anhydrous conditions. Synthesis was performed under agitation (180 rpm) using a magnetic hotplate stirrer fitted with an aluminum heat block at the specified temperature. At the specified endpoint, polymerization was quenched through exposure to atmospheric air, concentrated, and divided for crude analysis or subsequent extraction. Extraction was performed by solubilization in minimal DCM over 1 hr, followed by dropwise addition to cold IPA solvent (22.5mL IPA mL^−1^ DCM) on ice. Extracted precipitate was collected following decantation of the IPA and DCM mixture, dried by rotary evaporation (35mbar, 45℃, 45 min) and stored at 4℃ for subsequent analysis. Prior to thermal analysis and mass loss experiments, extracted polyesters were flash frozen in liquid nitrogen and lyophilized (0.200 mbar, −103℃, 18 hr).

**Table 1:**
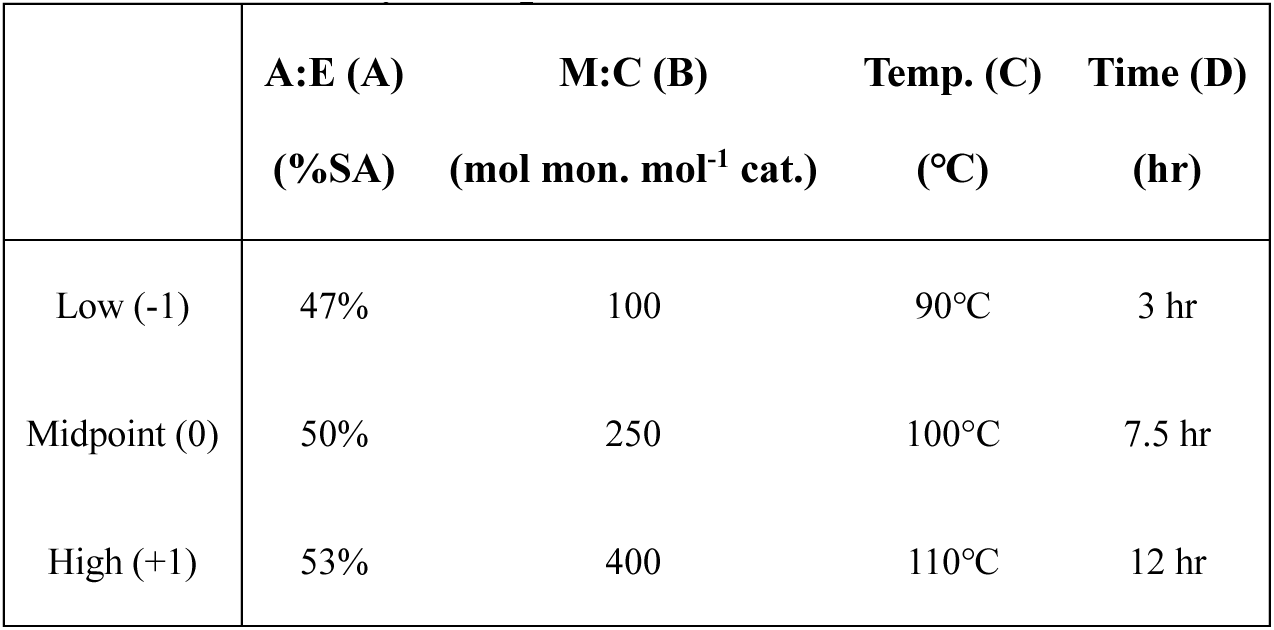
Summary of experimental conditions used in CCD.

Synthesis runs of PPS and PCS were conducted with variation of A:E ratio, M:C ratio, reaction temperature, and reaction time in a CCD design (*α*=1, face-centered axial points) implemented using StatEase DesignExpert Software (v.22.0.3) (Minneapolis, MN, United States) (**Table 1**). Total monomer concentration was held constant (0.01 mol mL ^−1^). Runs were completed sequentially in accordance with the random order dictated by StatEase DesignExpert (PCS: 28 total runs, PPS: 29 total runs) **(Table 2**). To assess system error, four centre points were included in the CCD. All SA and CHO used in synthesis were purified (by sublimation or distillation, respectively) in a single batch. PCS synthesis was conducted in one block of 28 runs. PPS synthesis was blocked (Block 1: runs 1-19, Block 2: runs 20-29) due to the need for two batches of PO distillation purification, and an additional run was included to evaluate block effects.

**Table 2:** Summary of input synthesis factor levels and material properties for succinate-based polyesters. A:E ratio: −1 = 47%, 0 = 50% +1 = 53% SA of total monomer; M:C ratio: −1 = 100, 0 = 250, +1 = 400 mol monomer mol^−1^ catalyst; T: −1 = 90℃, 0 = 100°C, +1 = 110℃; t: −1 = 3 hr, 0 = 7.5 hr, +1 = 12 hr. ND: Not Determined

| Input Factor |  |  |  |  | Crude Properties |  |  | Extracted Properties |  |  |  |
| --- | --- | --- | --- | --- | --- | --- | --- | --- | --- | --- | --- |
| Run | A:E<br>(A) | M:C<br>(B) | Temp.<br>(C) | Time<br>(D) | $\bar{M}_n$<br>(g mol<sup>-1</sup>) | PDI | Ester<br>Bond (%) | $\bar{M}_n$<br>(g mol<sup>-1</sup>) | PDI | Ester<br>Bond (%) | T<sub>g</sub><br>(°C) |
| PCS |  |  |  |  |  |  |  |  |  |  |  |
| 1 | 0 | 0 | 0 | 0 | 2592 | 1.32 | 93.9 | 2459 | 1.33 | 93.46 | 24.74 |
| 2 | 1 | 1 | -1 | -1 | 1612 | 1.24 | 86.21 | 1312 | 1.17 | 92.59 | ND |
| 3 | -1 | -1 | -1 | -1 | 2173 | 1.46 | 92.59 | 3108 | 1.54 | 93.46 | 13.68 |
| 4 | 1 | 1 | 1 | 1 | 2223 | 1.29 | 97.09 | 2634 | 1.34 | 95.24 | 13.99 |
| 5 | 1 | 1 | 1 | -1 | 1364 | 1.24 | 81.30 | 1578 | 1.57 | 87.34 | -43.21 |
| 6 | -1 | 1 | 1 | 1 | 2720 | 1.57 | 87.34 | 2779 | 1.57 | 88.5 | 22.21 |
| 7 | 0 | 0 | 0 | 0 | 2204 | 1.30 | 94.34 | 2463 | 1.30 | 94.34 | 24.99 |
| 8 | 1 | -1 | 1 | -1 | 1863 | 1.24 | 98.04 | 1822 | 1.26 | 98.04 | 31.38 |
| 9 | 1 | -1 | -1 | 1 | 1965 | 1.24 | 100.0 | 2425 | 1.22 | 100 | 28.20 |
| 10 | -1 | -1 | 1 | 1 | 2032 | 1.77 | 90.09 | 3213 | 1.77 | 93.9 | 25.05 |
| 11 | 1 | -1 | -1 | -1 | 1883 | 1.21 | 98.04 | 2082 | 1.21 | 98.52 | 25.72 |
| 12 | -1 | 1 | 1 | -1 | 1331 | 1.29 | 83.33 | 2434 | 1.47 | 88.11 | -34.13 |
| 13 | -1 | 1 | -1 | -1 | 1241 | 1.24 | 82.99 | 1452 | 1.34 | 70.92 | -57.85 |
| 14 | 1 | -1 | 1 | 1 | 1637 | 1.26 | 99.50 | 2002 | 1.30 | 99.01 | 19.20 |
| 15 | -1 | 1 | -1 | 1 | 1787 | 1.29 | 88.89 | 2780 | 1.31 | 94.34 | -5.17 |
| 16 | -1 | -1 | -1 | 1 | 2366 | 1.85 | 92.59 | 3960 | 1.68 | 94.79 | 20.13 |
| 17 | 1 | 1 | -1 | 1 | 1959 | 1.29 | 90.09 | 2969 | 1.31 | 92.59 | 8.62 |
| 18 | -1 | -1 | 1 | -1 | 2197 | 1.59 | 90.50 | 3020 | 1.76 | 92.59 | 33.27 |
| 19 | 0 | 0 | 0 | 0 | 2170 | 1.30 | 95.24 | 3475 | 1.37 | 94.79 | 31.00 |
| 20 | 0 | -1 | 0 | 0 | 1942 | 1.34 | 93.46 | 2520 | 1.37 | 93.9 | 23.11 |
| 21 | -1 | 0 | 0 | 0 | 2829 | 1.44 | 90.09 | 3139 | 1.52 | 91.74 | 25.57 |
| 22 | 0 | 0 | 0 | -1 | 1345 | 1.24 | 88.89 | 2131 | 1.32 | 93.9 | -11.60 |
| 23 | 0 | 0 | -1 | 0 | 2050 | 1.26 | 95.24 | 2563 | 1.30 | 95.69 | 17.85 |
| 24 | 0 | 0 | 1 | 0 | 2453 | 1.38 | 92.59 | 2684 | 1.43 | 93.46 | 16.92 |
| 25 | 0 | 0 | 0 | 1 | 2315 | 1.33 | 93.02 | 2842 | 1.41 | 94.34 | 18.37 |
| 26 | 1 | 0 | 0 | 0 | 2244 | 1.27 | 97.09 | 2463 | 1.32 | 96.62 | 13.71 |
| 27 | 0 | 1 | 0 | 0 | 1735 | 1.26 | 90.09 | 2363 | 1.41 | 90.5 | 6.55 |
| 28 | 0 | 0 | 0 | 0 | 2160 | 1.32 | 94.34 | 2825 | 1.33 | 95.24 | 23.16 |
| **PPS** |  |  |  |  |  |  |  |  |  |  |  |
| 1 | 0 | 0 | 0 | 0 | 4115 | 1.45 | 96.62 | 4668 | 1.49 | 97.09 | -23.40 |
| 2 | 1 | 1 | -1 | -1 | 2571 | 1.26 | 97.56 | 3160 | 1.26 | 98.04 | -21.32 |
| 3 | -1 | -1 | -1 | -1 | 3907 | 1.92 | 84.75 | 4706 | 1.75 | 85.11 | -12.49 |
| 4 | 1 | 1 | 1 | 1 | 3795 | 1.34 | 99.5 | 3811 | 1.36 | 99.5 | -24.57 |
| 5 | 1 | 1 | 1 | -1 | 2241 | 1.26 | 98.04 | 2514 | 1.25 | 98.04 | -22.13 |
| 6 | -1 | 1 | 1 | 1 | 4531 | 1.69 | 96.15 | 5183 | 1.74 | 96.15 | -22.87 |
| 7 | 0 | 0 | 0 | 0 | 4476 | 1.50 | 95.69 | 5279 | 1.59 | 95.69 | -19.78 |
| 8 | 1 | -1 | 1 | -1 | 2931 | 1.29 | 96.15 | 3002 | 1.31 | 96.15 | -22.79 |
| 9 | 1 | -1 | -1 | 1 | 2976 | 1.25 | 96.15 | 3405 | 1.26 | 96.15 | -20.90 |
| 10 | -1 | -1 | 1 | 1 | 3903 | 1.98 | 89.69 | 4586 | 1.74 | 90.5 | -16.18 |
| 11 | 1 | -1 | -1 | -1 | 3007 | 1.23 | 96.15 | 3415 | 1.25 | 96.15 | -22.49 |
| 12 | -1 | 1 | 1 | -1 | 2755 | 1.32 | 98.52 | 3345 | 1.30 | 98.52 | -22.74 |
| 13 | -1 | 1 | -1 | -1 | 2298 | 1.22 | 97.56 | 2483 | 1.20 | 98.04 | -25.09 |
| 14 | 1 | -1 | 1 | 1 | 2534 | 1.29 | 92.59 | 3035 | 1.33 | 92.17 | -26.88 |
| 15 | -1 | 1 | -1 | 1 | 5227 | 1.29 | 99.5 | 5153 | 1.37 | 99.5 | -22.23 |
| 16 | -1 | -1 | -1 | 1 | 4086 | 1.81 | 90.5 | 4790 | 1.85 | 91.74 | -10.57 |
| 17 | 1 | 1 | -1 | 1 | 3223 | 1.29 | 98.04 | 3778 | 1.30 | 98.52 | -21.95 |
| 18 | -1 | -1 | 1 | -1 | 3421 | 2.02 | 90.91 | 4454 | 1.93 | 93.02 | -17.69 |
| 19 | 0 | 0 | 0 | 0 | 3768 | 1.30 | 98.04 | 4341 | 1.39 | 98.04 | -23.21 |
| 20 | 0 | 0 | 0 | 0 | 5587 | 1.71 | 95.69 | 5647 | 1.84 | 96.15 | -18.81 |
| 21 | 0 | -1 | 0 | 0 | 3704 | 1.46 | 93.9 | 3756 | 1.50 | 93.9 | -21.14 |
| 22 | -1 | 0 | 0 | 0 | 5611 | 1.92 | 96.15 | 5660 | 1.97 | 97.09 | -19.55 |
| 23 | 0 | 0 | 0 | -1 | 3825 | 1.30 | 98.04 | 4046 | 1.28 | 98.04 | -24.13 |
| 24 | 0 | 0 | -1 | 0 | 4761 | 1.28 | 98.04 | 4926 | 1.31 | 98.04 | -20.52 |
| 25 | 0 | 0 | 1 | 0 | 4364 | 1.58 | 96.15 | 4583 | 1.56 | 96.15 | -22.71 |
| 26 | 0 | 0 | 0 | 1 | 4872 | 1.49 | 96.62 | 5058 | 1.54 | 96.15 | -20.68 |
| 27 | 1 | 0 | 0 | 0 | 3810 | 1.29 | 98.04 | 4039 | 1.49 | 98.04 | -24.07 |
| 28 | 0 | 1 | 0 | 0 | 3831 | 1.30 | 98.52 | 4023 | 1.26 | 98.52 | -19.75 |
| 29 | 0 | 0 | 0 | 0 | 4601 | 1.35 | 97.56 | 4820 | 1.48 | 97.56 | -22.20 |

### 2.3 Backbone Structure and Composition

Polymer backbone structure was characterized with proton nuclear magnetic resonance spectroscopy (^1^H-NMR). Samples were dissolved in CdCl_3_ (∼7 mg mL^−1^), filtered through a Kimwipe, then loaded into clean Wilmad precision tubes (200MHz frequency) at a minimum height of 40 mm. Analysis was performed using an AVANCE NEO 400MHz instrument (Bruker, Billerica, MA) using a basic 1D-1H experimental approach with 64 scans, followed by spectral analysis using MestreNova Software (v.15.0.1) (Mestrelab Research, Santiago de Compostela, Spain). All spectra were processed through an automated template: samples were linear phase shifted and referenced against the CdCl_3_ peak (7.26 ppm), automatic phase correction was performed followed by a Whittaker smoother baseline correction, and finally, automatic peak picking with specified integration regions in accordance with polymer structures and residual reagents was conducted (**Figure S1, Table S1**). Integrated spectra were subsequently analyzed for the percentage of ester bonds compared to ether bonds using Eqn 1 (PCS) or Eqn 2 (PPS).

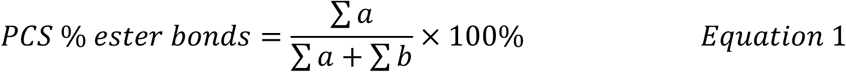

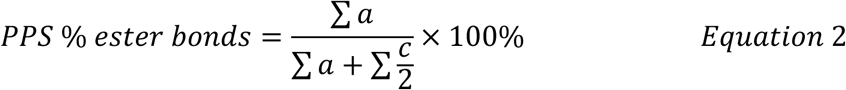

### 2.4 Molecular Weight Characterization

Polymer molecular weight was characterized by gel permeation chromatography (GPC) using an Agilent 1260 Infinity II GPC/SEC System equipped with two 300mm × 7.5mm PLgel mixed pore size columns (200 to 400,000 g mol^−1^) and a refractive index detector (Agilent Technologies, Santa Clara, United States). Polystyrene standards (peak molecular weight, M_p_ = 28,440 g mol^−1^, PDI =1.02) for system and column (n=12, M_p_ = 162 to 51,950 g mol^−1^) calibrations were used. Samples were prepared gravimetrically in THF (5mg mL^−1^), filtered (0.22 µm, polytetrafluoroethylene), and injected (50.00 µL) into the system (1.0mL min^−1^, 50℃). Conventional molecular weight analysis to determine *M̅_n_* and PDI was performed with Agilent GPC/SEC Software (v.2.8) (Agilent Technologies, Santa Clara, United States). Regions after 17.7 (PCS) and 17.25 (PPS) minutes (∼600 g mol^−1^, corresponding to monomer/catalyst complex) were omitted from integration and analysis as they were assumed to be residual monomer (**Figure S2**).

### 2.5 Glass Transition Temperature

Lyophilized extracted polyesters were analysed for T_g_ using a Discovery DSC 2500 (TA Instruments, New Castle, DE, United States) verified for performance using an indium standard and a polystyrene standard (*M̅_n_* = 2000 g mol^−1^). Samples (5 mg) were sealed in a Tzero aluminum pan (TA Instruments, New Castle, DE, United States) and analysed using a heat-cool-heat cycle. Samples were equilibrated to −80℃, heated to a melt (20℃ min^−1^ to 80℃) to erase previous thermal history, quenched (−10℃ min^−1^ to −80℃), and heated again (20℃ min^−1^ to 80℃). The second heat curve was used for T_g_ analysis by half tangent height through TRIOS Software (v.5.1.1) (TA Instruments, New Castle, DE, United States).

### 2.6 Mass Loss

Lyophilized extracted polyesters were subjected to base-catalyzed (NaOH, PCS: 1.0 M; NaOH, PPS: 1.0 M and 0.125 M) or neutral (phosphate buffer, 10 mM) hydrolysis as described elsewhere^32,33^. Polyester (40 mg) was added to a pre-weighed polypropylene conical tube (50 mL) and 10 mL of degradation solution was added. Samples were incubated for 7 d (37℃, 125 rpm) under base-catalyzed conditions, or assessed temporally under neutral conditions (1, 7, 14, 30 and 60 d). At experimental endpoint, samples were centrifuged (2500g, 5 min), supernatant was aspirated, and remaining polymer was rinsed three times with distilled water. Rinsed samples were lyophilized (0.220 mbar, 18 h) and weighed to yield a final mass. All masses were taken three times and averaged to ensure consistency, and mass loss was calculated as a relative change from initial mass across technical replicates (**Eqn. 3**).

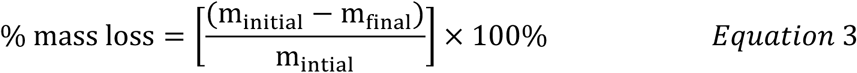

### 2.7 CCD Factor-Response Analysis & Validation

Material responses (*M̅_n_*, PDI, and % ester bonds for crude material; *M̅_n_*, PDI, % ester bonds, and T_g_ for extracted polyesters) were analysed in DesignExpert software using multi-linear regression on non-transformed data. Runs with extracted polyesters that did not yield sufficient polymer post-extraction were excluded from regression analysis of extracted properties. For each response variable, a quadratic model was fitted and analyzed using ANOVA. Terms with significant p-values (*α* = 0.05) (e.g., main effects, interactions, and quadratic terms) were retained, along with non-significant main effects linked to significant higher-order terms, to preserve the model hierarchy for the final reduced model.

Crude response models were validated using DesignExpert numerical optimization and confirmation tools. Using the numerical optimization tool, run parameters were generated that maximized crude *M̅_n_* (importance = 5), minimized crude PDI (importance = 5), and maximized crude ester bonds (importance = 3) for both PCS and PPS. A single synthesis point was selected for each PPS and PCS (**Table 3**), and three synthesis replicates were completed. Crude materials were then characterized following previous procedures for *M̅_n_*, PDI and ester bond % according to proximity to the predicted response value and the 95% CI around the predicted value.

**Table 3:** Experimental parameters used for validation of CCD models.

|  | A:E (A)<br>(%SA) | M:C (B)<br>(mol mon. mol<sup>-1</sup> cat.) | Temp. (C)<br>(°C) | Time (D)<br>(hr) |
| --- | --- | --- | --- | --- |
| PCS | 52.32 | 218.36 | 97.0 | 7.70 |
| PPS | 47.47 | 326.81 | 97.0 | 11.75 |

### 2.8 Cell Toxicity Assessment

Cell toxicity was assessed using an indirect toxicity test with serial dilution of conditioned media. PPS and PCS were degraded in DMEM (0.1 g mL^−1^, 37°C, 125 rpm) for 24h and 7d. At endpoints, degradation supernatant was collected and combined 1:1 (v:v) with DMEM containing 20% (v/v) FBS, and 2% (v/v) penicillin-streptomycin to generate degradation product conditioned cell culture media. Further dilutions were prepared in complete DMEM (10% FBS, 1% penicillin-streptomycin).

NIH 3T3 cells were treated (1.5×10^5^ cells cm^−2^, 24 hr) with a serial dilution of conditioned media, media only control, negative control, or positive control (10% DMSO). Following treatment, all wells were incubated with Hoechst 33342 (1:1000, DPBS), CFDA-SE (1:1000, DPBS), and PI (1:500, DPBS) for 10 min at 37°C, followed by imaging (4x magnification) using a BioTek Cytation 1 (Agilent Technologies, Santa Clara, CA) fitted with DAPI (447 nm), GFP (525 nm), and PI (647 nm) filter cubes. Cell counts were calculated using BioTek Gen5 software (v.2.07) (Agilent Technologies, Santa Clara, CA) with primary masking of the nucleus in the 447 nm channel. Dead cell counts were calculated using the same primary masking with secondary masking in the 647 nm channel constrained to the primary masks. Thresholds for positive staining were set based on the positive and negative controls. Cytotoxicity was calculated as percent dead using the dead count compared to the overall cell count.

### 2.9 Statistical Analysis

ANOVA analyses were used to developed factor-response models as described previously, using StatEase DesignExpert Software. Mass loss and cytotoxicity data were tested for normality and equality of variance using GraphPad Prism (v.10.0) (Dotmatics, Boston, MA); all data met the tested assumptions. Parametric one-way or two-way ANOVA followed by pairwise comparisons with post-hoc pairwise comparison tests were used to determine the statistical significance and assess the interactive effects of factors. Specific statistical tests for mass loss and cytotoxicity data are noted in the respective figure caption and/or results text.

## 3 RESULTS

### 3.1 Systematic ROCOP Synthesis and Characterization of SA Polyesters

Succinate-based polyester materials were synthesized using a ROCOP approach in solution, combining SA with PO or CHO in the presence of a Cr(III) salen /PPNCl catalyst/co-catalyst system to yield PPS and PCS (**Figure 1A**). To efficiently appreciate factor-response relationships, synthesis was performed using a CCD approach to vary A:E ratio (Factor A), M:C ratio (Factor B), reaction temperate (Factor C) and reaction time (Factor D), ensuring both linear and curved responses could be modeled effectively (**Table 1**)^34^. All synthesis runs successfully generated polyesters that exhibited qualitative properties ranging from a brittle solid to viscous liquid at physiologic conditions (37°C, pH 7.4), with structure confirmed by ^1^H-NMR analysis (representative spectra: **Figure 1B**). Synthesized polyesters were characterized for *M̅_n_*, PDI, and the ester bond % on both crude and extracted polyesters, as well as T_g_ following extraction (**Table 2)**.

**Figure 1:**
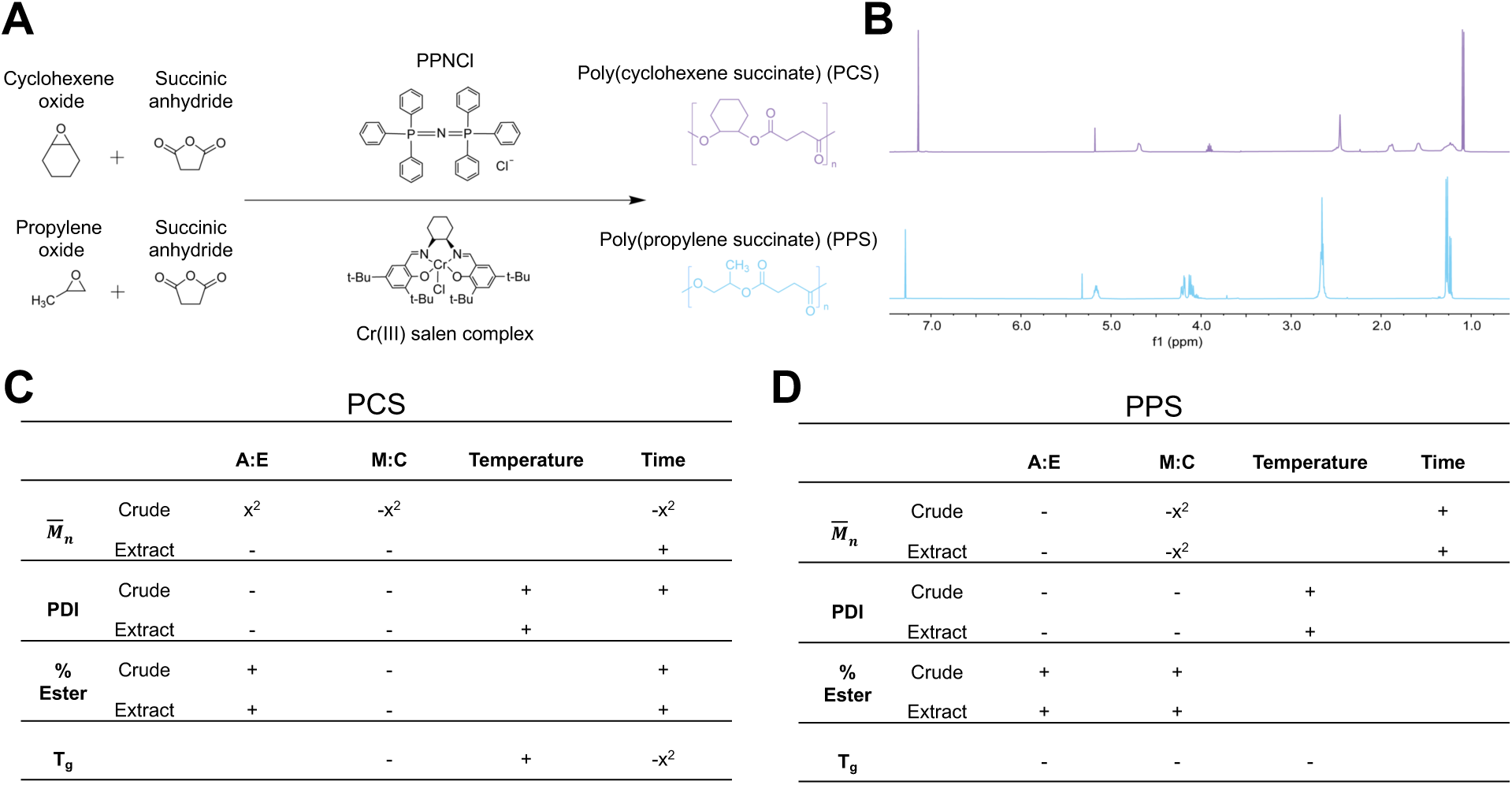
ROCOP synthesis and characterization of SA based polyesters. (A) SA was combined with CHO or PO to generate PCS and PPS polyesters in solution with a Cr(III) salen/PPNCl catalyst/co-catalyst in solution. (B) Representative ^1^H NMR of PCS (purple, top) and PPS (blue, bottom) polyesters. (C-D) Summary of significant synthesis factor – material response relationships for (C) PCS and (D) PPS materials from multi-linear regression of CCD. + or – indicate linear relationship is dominant. +x^2^ or -x^2^ indicate quadratic relationship is dominant. No symbol indicates non-significant.

Factor-response relationships were evident in the diversity of the material properties observed for PCS (*M̅_n_*: 1241 to 2829 g mol^−1^; PDI: 1.213 to 1.848; ester bonding: 81.3 to 100%; T_g_ :−55.85 to 33.27 ℃) and PPS (*M̅_n_*: 2241 to 5611 g mol^−1^; PDI: 1.223 to 2.023; ester bonding: 84.75 to 99.5%; T_g_ :−26.88 to −10.57℃). PPS had a notably higher *M̅_n_* across all samples, which is further emphasized when considering the degree of polymerization (DP) (mean DP: PCS: 13, PPS: 24) given PO (50.08 g mol^−1^) has a lower molar mass then CHO (98.14 g mol^−1^). Material property responses were subsequently related to input factors using empirical regression models, yielding significant positive (+) or negative (−) linear (x) or quadratic (x^2^) relationships (**Figure 1C,D**). All synthesis factors had significant relationships to material properties; A:E and M:C ratios were significant (p<0.05) to all material properties, except for A:E ratio to PPS T_g_, highlighting the importance of feed ratios. Reaction conditions also demonstrated significance to assessed material properties; temperature significantly impacted PDI and T_g_ for both polyesters, and reaction time was significant to PCS and PPS *M̅_n_*, PCS T_g_, PCS ester bond percentage, and PCS crude PDI. The magnitude and interplay of synthesis factors was further assessed through developed regression models, described on a property-specific basis in the following sections.

Polyester extraction posed challenges in factor-response models, wherein low degree of polymerization and excess SA monomer complicated recovery yield in solvent extraction. Polyesters with excessive amounts of residual SA required excess DCM addition for complete solubilization of crude material, which subsequently increased the polymer solubility in the extraction mixture (IPA) and reduced precipitation. Peak integration quantification of extracted PCS polyesters from runs 2, 5, 12, 13, 15 and 17 showed excess SA (SA : ester >1) remaining in the extracted polyester; this was particularly evident in Run 2 (SA : ester = 648.24), which did not present a detectible T_g_, suggesting limited polymer content. For all remaining materials, the relative amount of SA was decreased during polyester extraction. Excess SA in PPS polyesters was minimal overall compared to PCS polyesters, save for PPS runs 2 and 5 which showed excess residual SA (SA : ester >2) following extraction. Epoxide was not detected in significant excess (CHO/ester >1, PO/ester >2) for any PCS or PPS samples, likely due to its volatility. *M̅_n_* increased for all materials between crude and extracted polymer analysis, except for PCS runs 1, 2, and 8 and PPS run 15, whereas PDI and the ester bond % were well conserved.

### 3.2 Factor-Material Response Relationships

#### 3.2.1 Molecular Weight

Synthesis factors exhibited significant relationships to crude and extracted polymer *M̅_n_* (ANOVA Tables: **Table S2-S9**). Molecular weight analysis on crude materials was performed following exclusion of low mass molecules evident in elution traces (**Figure S2**). A reduced model for crude *M̅_n_*, generated after removal of non-significant factors from the full ANOVA analysis, was significantly related to synthesis factors for PCS (p = 0.0002; **Eqn. 4**, **Figure 2A**) and PPS (p<0.0001; **Eqn. 5**, **Figure 2B**).

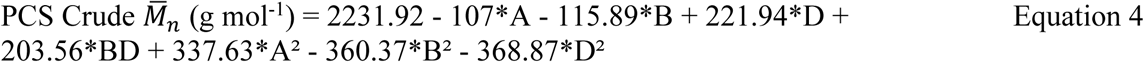

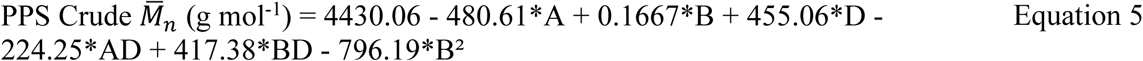

**Figure 2.**
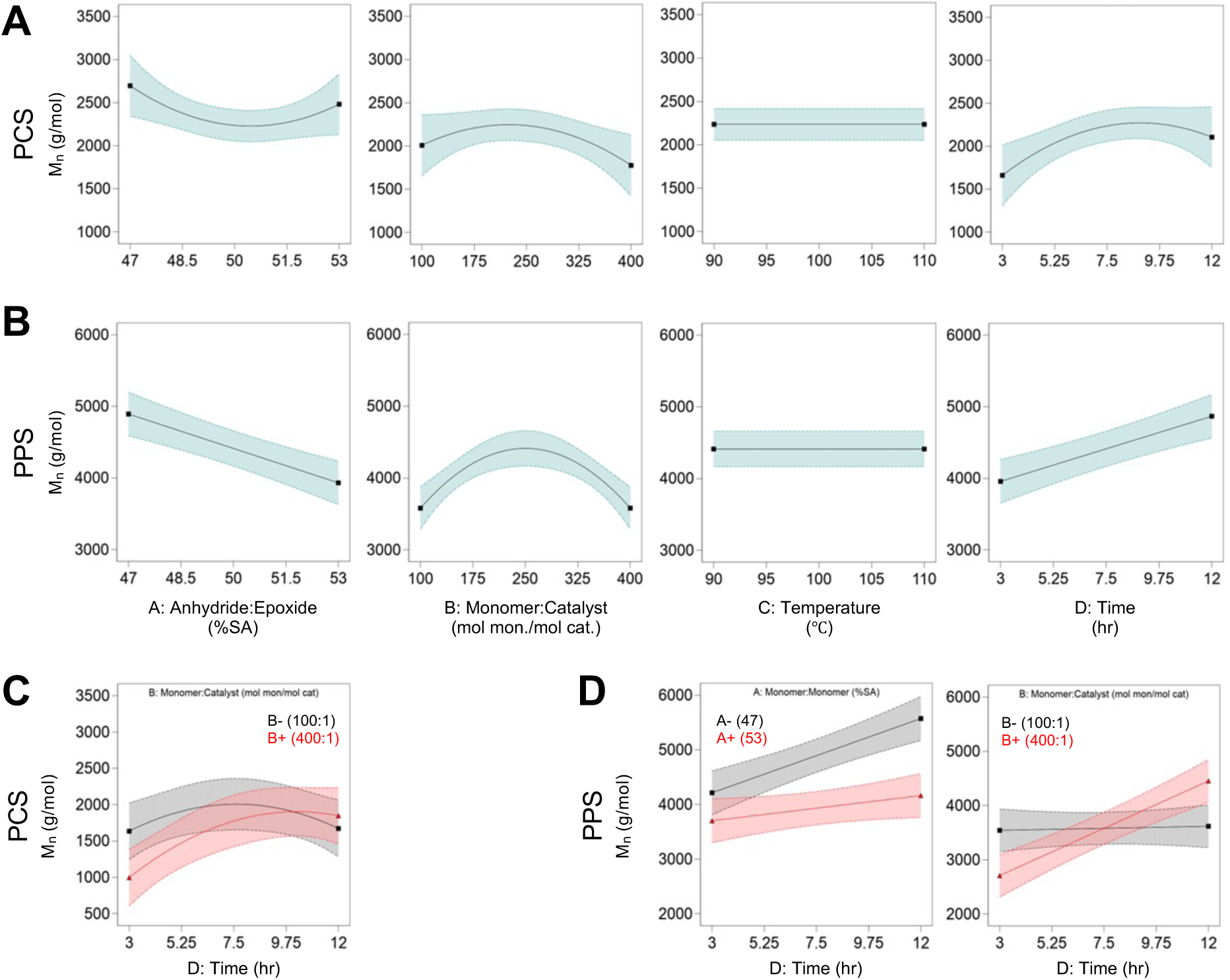
Factor-response models highlight relationships between synthesis variables and crude polymer *M̅_n_*. (A-B) Visual representation of expected (A) PCS and (B) PPS crude *M̅_n_* as a function of synthesis factors. Data was empirically modelled and is presented within the high and low experimental bounds of each factor. Curves represent predictions from the reduced regression model shown across the high (+1) and low (−1) bounds of each factor while holding other factors at their midpoint (0). (C-D) Interaction plots illustrate the influence of significant interaction effects for (C) PCS and (D) PPS. Each plot shows two curves across the experimental range of the x-axis factor, with the interacting factor fixed either at its high (red) or low (grey) level. All other factors were held constant at their midpoint (coded value: 0). Shaded regions in all plots denote 95% confidence intervals.

Both materials investigated exhibited significant linear and quadratic relationships to feed ratios. The quadratic effects of feed ratios had a stronger influence on *M̅_n_* than their linear effects in PCS (A:E and M:C ratios) and PPS (M:C ratio) materials, observed visually through a quadratic minimum for the A:E ratio and maximum for the M:C ratio. A linear relationship to reaction time was also observed, with an increase in *M̅_n_* over time. PCS materials also demonstrated a significant quadratic relationship, but this was less certain when considering the breadth of the 95% CI that may suggest a plateau at 9 hours.

A CCD approach to factor-response analysis enabled a quantitative empirical appreciation of the interaction effects of input factors on material properties. M:C ratio and time had significant interaction effects; PCS with high (+1) M:C ratio had a lower *M̅_n_* at 3 hours and showed a positive increase in *M̅_n_* over time in PCS materials (**Figure 2C**), whereas PPS *M̅_n_* was only notably impacted by time at high (+1) M:C ratio (**Figure 2D**). Time also exhibited an interaction effect with the A:E ratio in PPS materials, with a stronger influence on *M̅_n_* at a low (−1) A:E ratio (**Figure 2D**).

The reduced models for the *M̅_n_* of extracted polyesters were comparably significantly related to synthesis factors for PCS (p <0.0001, **Eqn. S1, Figure S3A**) and PPS (p<0.0001; **Eqn. S2, Figure S3B**). A:E ratio, M:C ratio, and time factors had significant linear relationships with extract *M̅_n_* comparable to the crude materials. Quadratic relationships were less impactful in extracted materials, except for M:C ratio in PPS, which exhibited a notable maximum at the midpoint of the range assessed. The interaction of M:C ratio and time remained significant for both formulations (**Figure S3C, D)**. The interaction of A:E ratio and M:C ratio was uniquely significant for extracted *M̅_n_* of PCS, with a notable negative impact with increasing SA feed at low (−1) M:C ratio (**Figure** S3C).

#### 3.2.2 Polydispersity Index

A low PDI is a particularly advantageous property of polyesters developed by ROCOP approaches^24,35^ and has an appreciated impact on degradation properties of polyesters^36–38^. Comparable to *M̅_n_* assessment, crude and extracted PDI exhibited significant relationships to synthesis factors (ANOVA Tables: **Table S10-S17**). Significant reduced models for crude PDI for PCS (p<0.0001; **Eqn. 6**, **Figure 3A**) and PPS (p<0.0001; **Eqn. 7**, **Figure 3B**) were determined.

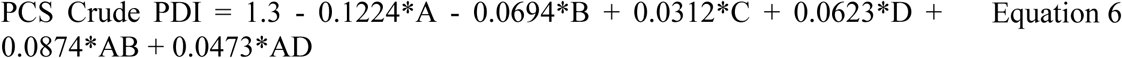

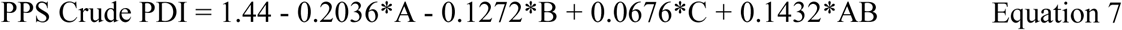

**Figure 3.**
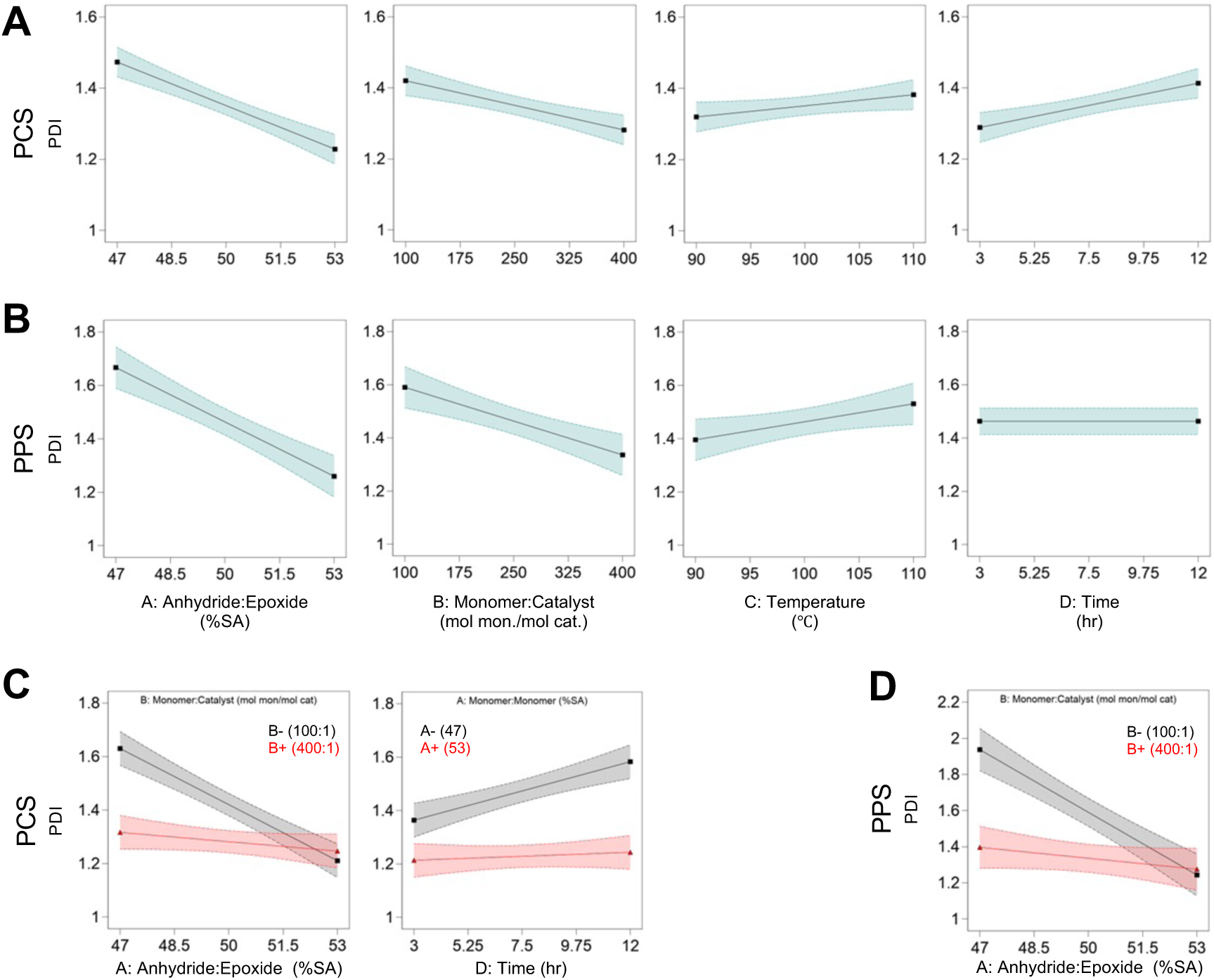
Factor-response models highlight relationships between synthesis variables and crude polymer PDI. (A-B) Visual representation of expected (A) PCS and (B) PPS crude PDI as a function of synthesis factors. Data was empirically modelled and is presented within the high and low experimental bounds of each factor. Curves represent predictions from the reduced regression model shown across the high (+1) and low (−1) bounds of each factor while holding other factors at their midpoint (0). (C-D) Interaction plots illustrate the influence of significant interaction effects for (C) PCS and (D) PPS. Each plot shows two curves across the experimental range of the x-axis factor, with the interacting factor fixed either at its high (red) or low (grey) level. All other factors were held constant at their midpoint (coded value: 0). Shaded regions in all plots denote 95% confidence intervals.

Crude PDI for PCS was significantly correlated to all synthesis factors, whereas time was excluded for PPS. These relationships were comparable for both polyesters; crude PDI decreases as the relative amount of SA fed increases and as the total amount of monomer increases relative to catalyst and increases with temperature (**Figure 3A, B**). PCS crude PDI also increases with time (**Figure 3A**). A:E ratio and M:C ratio had significant interaction effects for both polyesters; low M:C ratio polyesters had a higher PDI at low (−1) A:E ratio conditions compared to the high (+1) A:E ratio condition, whereas high M:C ratio polyesters did not show as much PDI variation with A:E ratio (**Figure 3C, D**). PCS polyesters also had significant interactions between A:E ratio and time; the low (−1) A:E ratio condition presented an increase in PDI with time, while the high (+1) A:E ratio condition showed a less prominent trend (**Figure 3C**).

PDI of extracted polyesters similarly demonstrated significant relationships to synthesis factors for PCS (p <0.0001, **Eqn. S3, Figure S4A**) and PPS (p<0.0001; **Eqn. S4, Figure S4B**), albeit with fewer variables contributing to the response. A:E and M:C ratios had a significant linear relationship with both polyesters, with comparable correlations to crude PDI (**Figure S4A, B**). Temperature was a significant positive linear term in the reduced model for PCS, but was non-significant for PPS. As with crude PDI, the A:E and M:C ratios presented a significant interaction term, with a stronger negative relationship between PDI and A:E ratio observed for low (−1) M:C ratio polyesters (**Figure S4C, D**).

#### 3.2.3 Alternating Structure

ROCOP synthesis can yield both ester and ether linkages resultant of double epoxide insertion^39^. Ester and ether bonds result in polyesters with different bulk material properties, where ether bonds are particularly less susceptible to hydrolysis than ester bonds^40^. To appreciate this, we quantified the percentage of polymer backbones that were ester linkages (% ester bond) through ^1^H NMR analysis as a measure of the degree of alternating structure. Crude and extracted % ester exhibited significant relationships to synthesis factors (ANOVA Tables: **Table S18-S25**), which are empirically modelled through reduced models for PCS (p<0.0001; **Eqn. 9**, **Figure 4A**) and PPS (p<0.0001; **Eqn. 10, Figure 4B**).

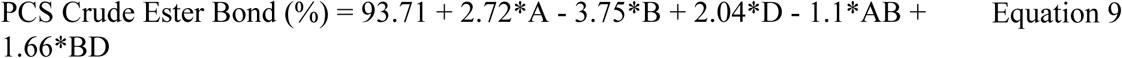

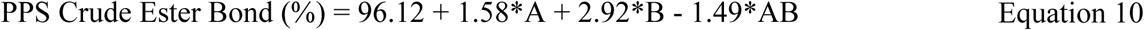

**Figure 4.**
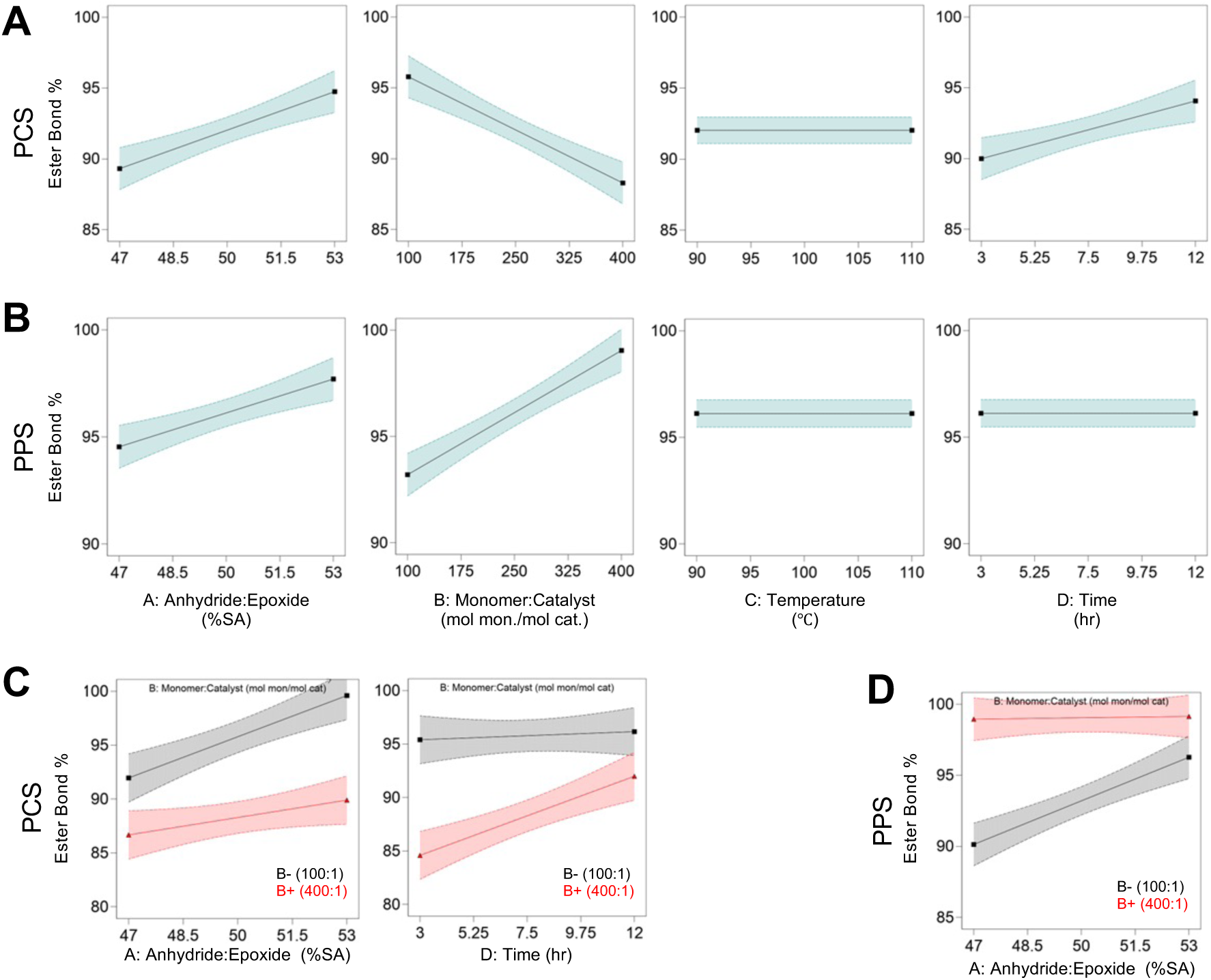
Factor-response models highlight relationships between synthesis variables and crude polymer alternating structure. (A-B) Visual representation of expected (A) PCS and (B) PPS crude ester bond frequency as a function of synthesis factors. Data was empirically modelled and is presented within the high and low experimental bounds of each factor. Curves represent predictions from the reduced regression model shown across the high (+1) and low (−1) bounds of each factor while holding other factors at their midpoint (0). (C-D) Interaction plots illustrate the influence of significant interaction effects for (C) PCS and (D) PPS. Each plot shows two curves across the experimental range of the x-axis factor, with the interacting factor fixed either at its high (red) or low (grey) level. All other factors were held constant at their midpoint (coded value: 0). Shaded regions in all plots denote 95% confidence intervals.

The percentage of ester bonds had a significant linear relationship to A:E ratio, showing an increase in alternating structures as the amount of fed anhydride increased relative to epoxide (**Figure 4A, B).** Interestingly, other model parameters differed in significance or correlative relationship between PCS and PPS. While both formulations were significantly impacted by M:C ratio, the alternating structure decreased as more monomer was fed relative to catalyst in PCS and increased for PPS. Frequency of ester bonding was positively correlated with time in crude PCS (**Figure 4A**). The interaction of A:E and M:C ratios was significant for both formulations; ester bonding increased with increasing A:E ratio for both materials, but PCS showed more ester bonding in the low (−1) M:C ratio condition (**Figure 4C)** in contrast to more relative ester bonding in the high (+1) M:C ratio condition in PPS materials (**Figure 4D**). PCS was further impacted by the interaction of M:C ratio and time; materials synthesized with high (+1) M:C ratio had lower ester bonding at the low (−1) time condition and increased ester bonding with time (**Figure 4C)**.

Alternating structure had comparable significant relationships to synthesis factors in extracted PCS (p <0.0001, **Eqn. S5, Figure S5A**) and PPS (p<0.0001; **Eqn. S6, Figure S5B**), with deviations from crude analysis in the observed interaction effects. The interaction between A:E and M:C ratios was maintained for PPS but was not significant for PCS. The alternating structure of extracted PPS was also significantly impacted by the interaction between temperature and time; low (−1) temperature conditions increased ester bonding with time, whereas high (+1) temperature conditions saw a decreasing linear relationship to ester bonding as time increased. However, the 95% CI in this case do overlap considerably (**Figure S5C**).

#### 3.2.4 Glass Transition Temperature

T_g_ is central to understanding chain entanglement and flexibility of a specific polymer, governing diffusion of water into and degradation products out of the polyester matrix^41^. Unlike other material property responses, T_g_ was only performed on extracted polyesters given the potential confounding impact of remaining solvent, monomer, or catalyst on analysis. T_g_ models showed significant relationships to synthesis factors (ANOVA Tables: **Table S26-S29**), represented through reduced models for PCS (p<0.0001; **Eqn. 11, Figure 5A**) and PPS (p<0.0001; **Eqn. 12, Figure 5B**).

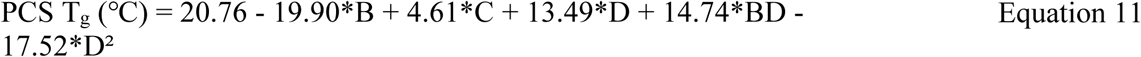

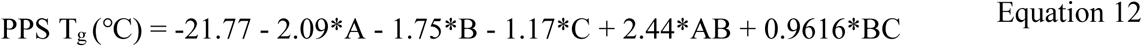

**Figure 5:**
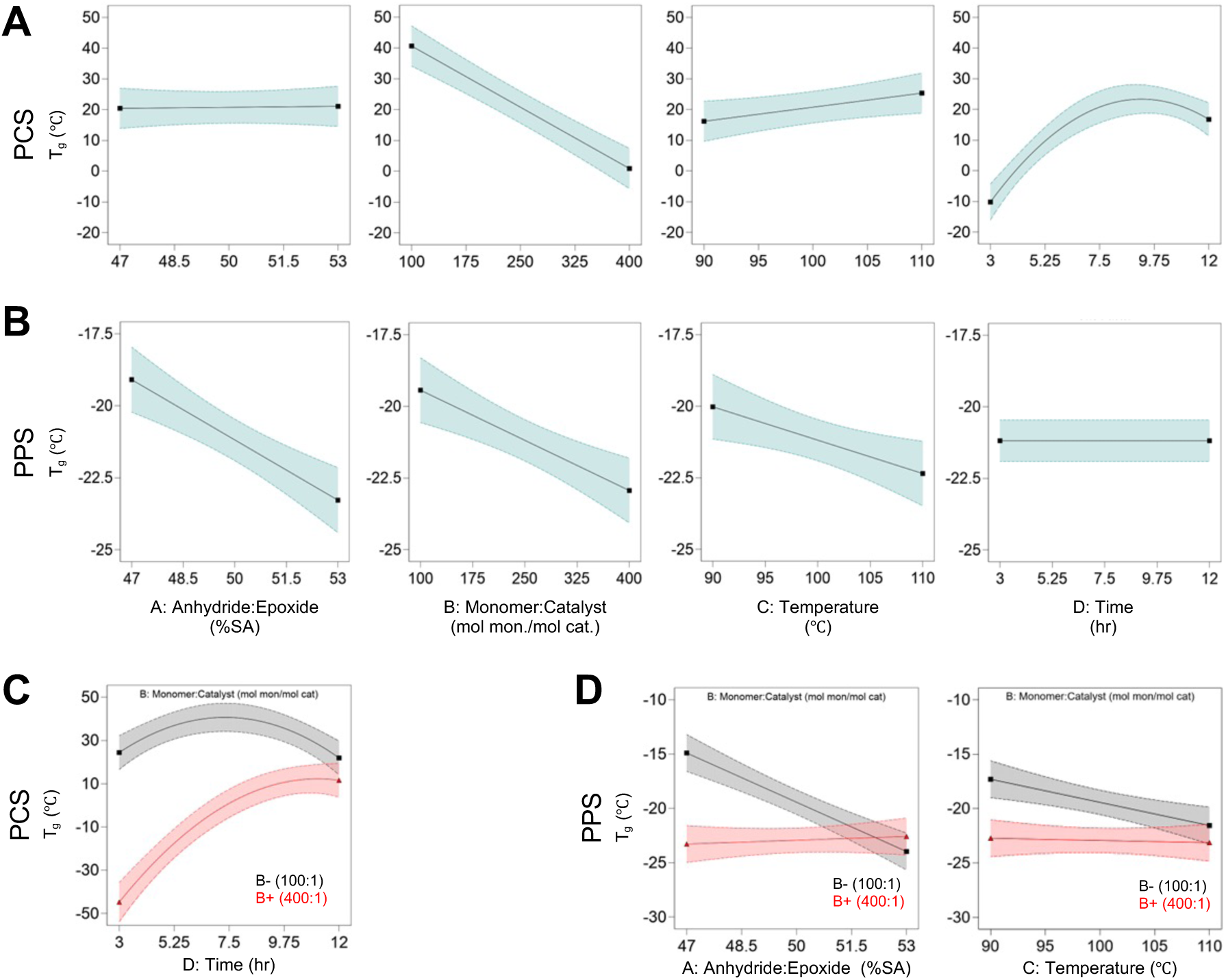
Factor-response models highlight relationships between synthesis variables and polyester T_g_. (A-B) Visual representation of expected (A) PCS and (B) PPS T_g_ as a function of synthesis factors. Data was empirically modelled and is presented within the high and low experimental bounds of each factor. Curves represent predictions from the reduced regression model shown across the high (+1) and low (−1) bounds of each factor while holding other factors at their midpoint (0). (C-D) Interaction plots illustrate the influence of significant interaction effects for (C) PCS and (D) PPS. Each plot shows two curves across the experimental range of the x-axis factor, with the interacting factor fixed either at its high (red) or low (grey) level. All other factors were held constant at their midpoint (coded value: 0). Shaded regions in all plots denote 95% confidence intervals.

PCS T_g_ decreased with increased M:C ratio, increased with polymerization temperature, and exhibited a general positive trend with polymerization time along with maxima behaviour (**Figure 5A**). Increases in M:C ratio similarly decreased T_g_ in PPS materials but exhibited a negative relationship with polymerization temperature and A:E ratio (**Figure 5B**). PPS materials also demonstrated a notably smaller range of T_g_ in the range of input factors assessed. M:C ratio played a prominent role in observed interaction effects; in PCS materials, high (+1) M:C condition had a lower T_g_ at low (−1) time conditions (and T_g_ increased with time), whereas low (−1) M:C had much higher T_g_ and showed slight quadratic behaviour over time (with a maximum T_g_ at the time mid-point) (**Figure 5C**). In PPS materials, M:C ratio had significant interaction effects with A:E ratio and temperature (**Figure 5D**).

#### 3.2.5 Model Validation

To understand model utility and validate the empirical models generated for PCS and PPS, we synthesized new polyesters with predicted optimized *M̅_n_*, PDI, and ester bonding to compare their material responses to model expectations. PCS validation synthesis demonstrated crude *M̅_n_*, PDI, and ester bonding within the 95% CI of the model predicted values (**Table 4**). The mean crude *M̅_n_* was within 64 g mol^−1^ and 40.15 g mol^−1^ from the predicted value for PCS and PPS, respectively. PDI was similarly well-predicted for PCS (Δ = 0.01) but did not fall within the 95% CI for PPS (Δ = 0.51). The crude ester bond percentage was well-predicted for both material formulations. In all scenarios, variation of the properties was minimal across the three technical replicates of synthesis.

**Table 4:**
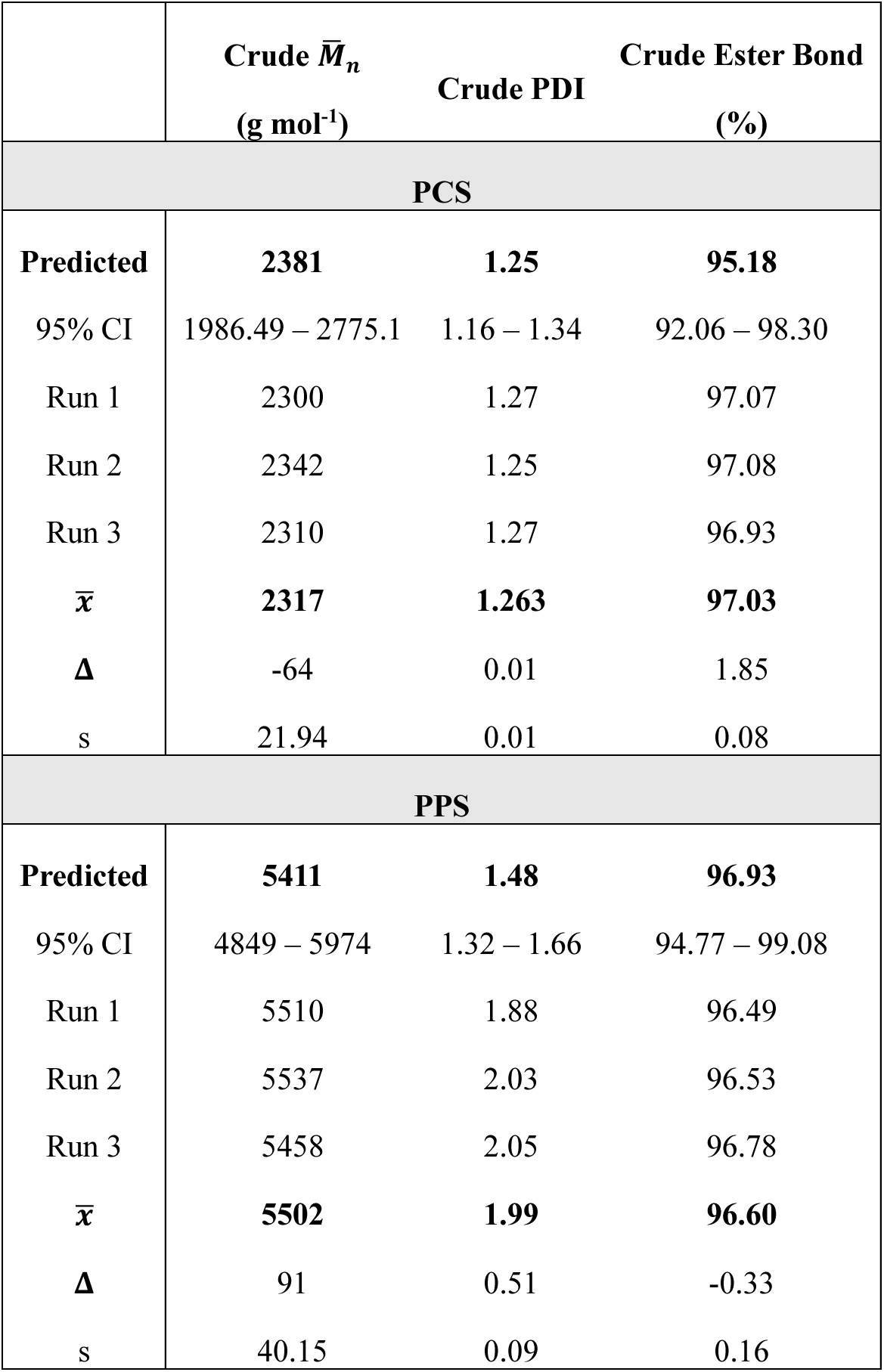
Validation of material factor-response models for maximized crude *M̅_n_* and alternating structure with minimized PDI. PCS synthesis factor levels: A:E (A): 52.32% SA, M:C (B): 218.36 mol monomer mol^−1^ catalyst, T (C): 97°C, t (D): 7.70 hr. PPS synthesis factor levels: A:E ratio (A): 47.47% SA, M:C ratio (B): 326.81 mol monomer mol^−1^ catalyst, T (C): 97°C, t (D): 11.75 hr. Synthesis was performed in triplicate. Δ = difference from predicted value

| | Crude $\bar{M}_n$<br>(g mol<sup>-1</sup>) | Crude PDI | Crude Ester Bond<br>(%) |
| --- | --- | --- | --- |
| **PCS** |  |  |  |
| **Predicted** | **2381** | **1.25** | **95.18** |
| 95% CI | 1986.49 – 2775.1 | 1.16 – 1.34 | 92.06 – 98.30 |
| Run 1 | 2300 | 1.27 | 97.07 |
| Run 2 | 2342 | 1.25 | 97.08 |
| Run 3 | 2310 | 1.27 | 96.93 |
| $\bar{x}$ | **2317** | **1.263** | **97.03** |
| $\Delta$ | -64 | 0.01 | 1.85 |
| s | 21.94 | 0.01 | 0.08 |
| **PPS** |  |  |  |
| **Predicted** | **5411** | **1.48** | **96.93** |
| 95% CI | 4849 – 5974 | 1.32 – 1.66 | 94.77 – 99.08 |
| Run 1 | 5510 | 1.88 | 96.49 |
| Run 2 | 5537 | 2.03 | 96.53 |
| Run 3 | 5458 | 2.05 | 96.78 |
| $\bar{x}$ | **5502** | **1.99** | **96.60** |
| $\Delta$ | 91 | 0.51 | -0.33 |
| s | 40.15 | 0.09 | 0.16 |

### 3.3 Mass Loss

Material property characterization was paired with degradation assessment of extracted PCS and PPS under accelerated alkaline conditions, enabling comparison of the impact of bond stability, alternating structure, and molecular weight on the rate of mass loss^42^. PPS materials (Run 7, **Table 2** exhibited near complete mass loss (94.6 ± 0.5 %) under accelerated conditions (1.0 M NaOH), which was notably different from PCS (6.7 ± 2.1 %) with the same synthesis factor inputs (**Figure 6A**). Further analysis of PPS and PCS material libraries were performed under unique accelerated conditions (7 days; PCS: 1.0 M NaOH, PPS: 0.125 M NaOH) to appreciate relative differences in mass loss for each formulation. Mass loss was highly variable under these conditions (PCS: 4.79 to 99.4%, PPS: 16.62 to 100%) and appeared to be related to extracted *M̅_n_* and PDI (**Figure 6B, C)**. Several materials exhibited complete mass loss over this timeline, especially in the PPS library, limiting the ability to perform statistical analysis with confidence. To appreciate how these findings translated to degradation relevant to biomaterial applications, we assessed mass loss of extracted polyesters synthesized with optimized *M̅_n_*, PDI, and ester bonding (**Table 4, Run 1 for both PCS and PPS**) under physiologically relevant conditions (10 mM phosphate buffer, 37°C) for up to 60 days. Mass loss increased temporally in both formulations and was significantly higher in PPS materials (Two-way ANOVA; Material: p<0.0001, Time: p<0.0001) (**Figure 6D**). Replicate synthesis runs exhibited a significant difference in mass loss at 60 days (One-way ANOVA; PCS: p=0.0412, PPS: p=0.0437), albeit differences were minor (**Figure 6E**). The temporal *M̅_n_* (**Figure 6F**) and PDI (**Figure 6G**) was unchanged in both formulations over the same timeline, save for a slight reduction in PPS PDI at 30 and 60 days from baseline.

**Figure 6:**
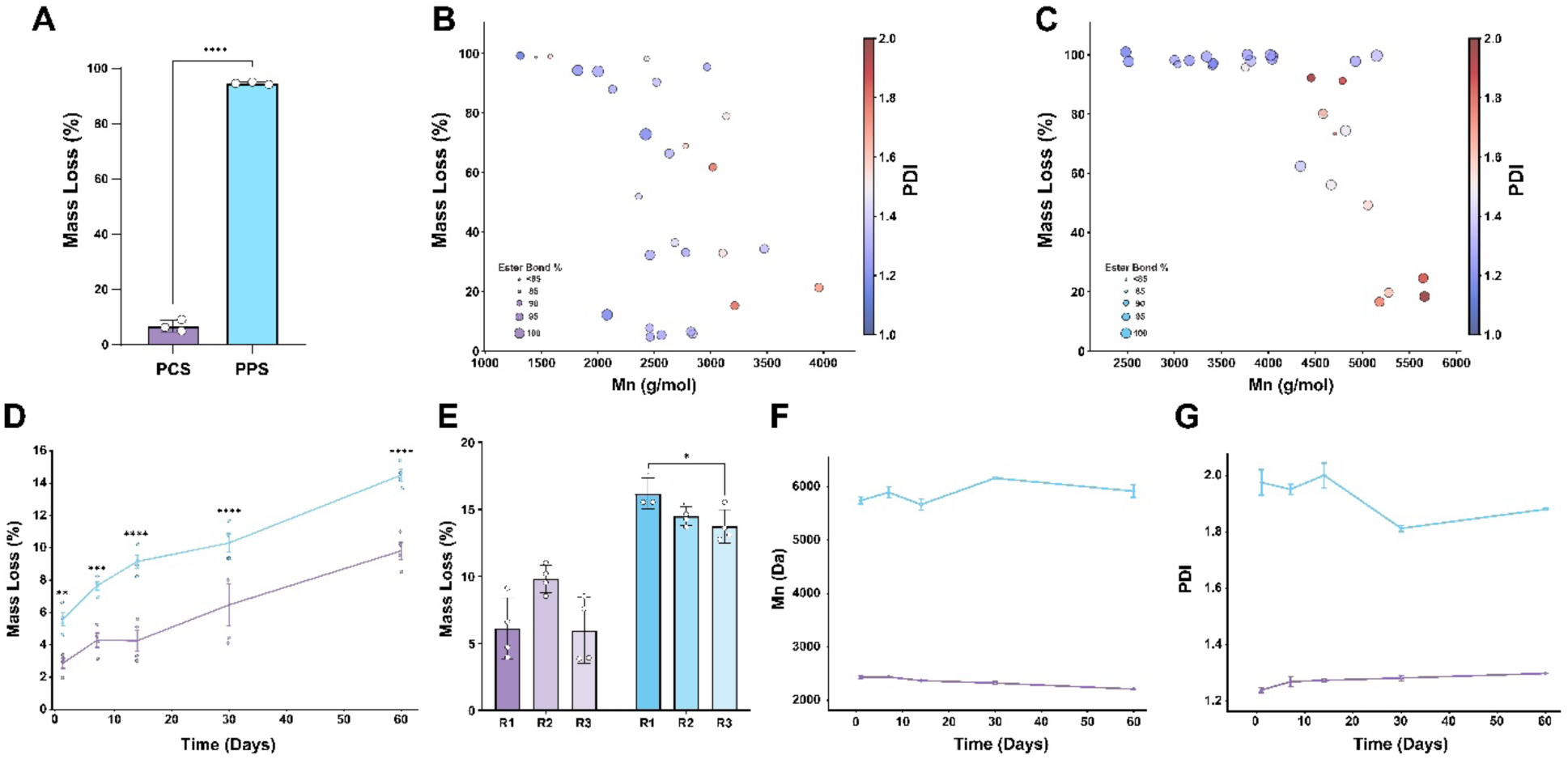
Hydrolytic degradation of PCS and PPS materials is composition-specific. (A) Accelerated hydrolysis (1.0 M NaOH, 7 days) of PCS and PPS materials with matched synthesis factors (Run 7). n = 3. Mean ± SD. Paired t-test. (B-C) Mass loss of extracted (B) PCS and (C) PPS under accelerated conditions (PCS: 1.0 M NaOH, PPS: 0.125 M NaOH, 7 days). Data presented as a function of extracted *M̅_n_* (x-axis), ester bond % (dot size), and PDI (heat map statistic). Each data point is presented as the mean of n=4 technical replicates from each synthesis run. (D-G) Temporal hydrolysis of PCS (purple) and PPS (blue) materials over 60 days in physiologically relevant conditions (37°C, 10 mM phosphate buffer). (D) Mass loss increases temporally over 60 days. n ≥ 3. Mean ± SD. Two-way ANOVA followed by Tukey’s multiple comparisons test. Significance indicated between materials at each time point. (E) Comparative mass loss between synthesis runs at 60 days. n ≥ 3. Mean ± SD. One-way ANOVA within each formulation followed by Tukey’s multiple comparisons test. (F) *M̅_n_* and (G) PDI are unchanged over 60 days. n =2. Mean ± SD. Significance indicated as ****p<0.0001, ***p<0.001, *p<0.05.

### 3.4 Cytotoxicity Assessment

To appreciate the potential applicability of PCS and PPS materials as degradable biomaterials, we assessed the toxicity of media conditioned with extracted polyesters synthesized with optimized *M̅_n_*, PDI, and ester bonding (**Table 4, Run 3**). Basal culture media was conditioned with 0.01 g polymer mL^−1^ for 24 h or 7 d prior to treatment of NIH3T3 fibroblasts. PCS and PPS conditioned media exhibited limited cytotoxic effects on NIH3T3 murine fibroblasts when treated in the diluted concentration range of 0.05 g polymer mL^−1^ (2X dilution of original conditioned media) to 0.001 g polymer mL^−1^ (100X dilution of original conditioned media). PCS had no observed cytotoxic effects across any concentrations (**Figure 7B, C**). PPS presented a significant reduction in viability at the highest concentration (0.05 g PPS mL^−1^), but the magnitude of toxicity observed (24 h: 2.09 ± 0.91%, 7 d: 1.73 ± 1.46%) was minimal and well within the acceptable range (**Figure 7D, E**).

**Figure 7.**
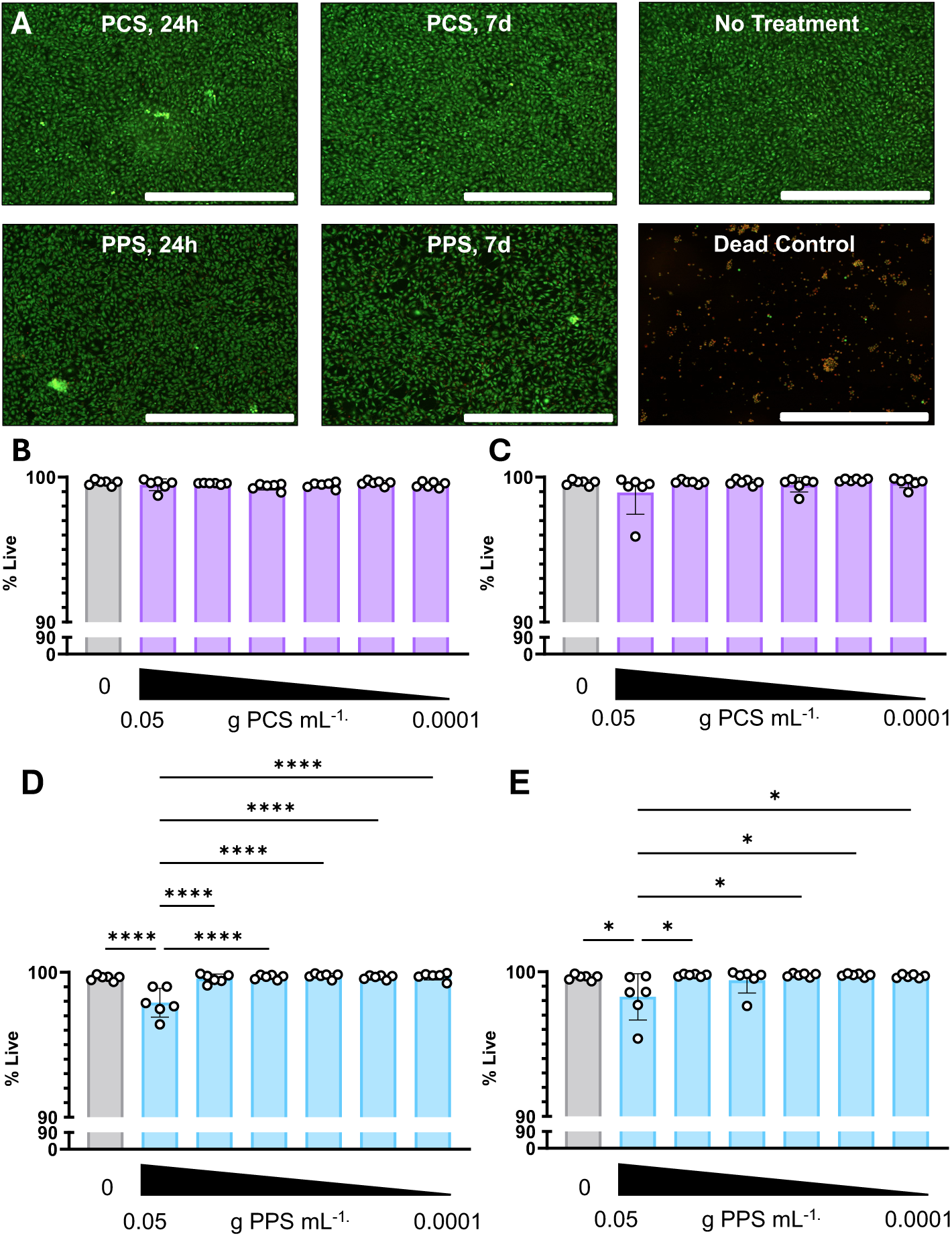
PCS and PPS degradation products are non-toxic. (A) Representative endpoint images (24 hr, 7 d; 0.05 g polymer mL^−1^) of NIH3T3 visually demonstrate high viability compared to untreated and positive (dead) control. Green: CFDA-SE (live). Red: PI (dead). Scale bar = 1000 μm. (B-E) Quantification of NIH3T3 cell viability treated with media pre-conditioned with degradable (B-C) PCS or (D-E) PPS for (B,D) 24 hr or (C,E) 7 d. Gradient represents decreasing concentration (left to right, serial 2X dilution) from 0.05 g polymer mL^−1^ to 0.001 g polymer mL^−1^. n = 6. Mean ± SD. One-way ANOVA followed by post-hoc Tukey’s multiple comparisons test. Significance indicated as ****p<0.0001, * p<0.05

## 4 DISCUSSION

The predictable development of degradable polyester biomaterials with diverse properties has been limited with traditional synthesis methodologies by broad PDI or low monomer diversity. Advances in ROCOP of cyclic anhydrides and epoxides have demonstrated potential for overcoming those limitations with improved monomer diversity and reproducible, precise material properties. Here, we investigated ROCOP for its utility in polyester biomaterial development through a CCD synthesis of PCS and PPS using a Cr(III) salen and PPNCl catalyst/cocatalyst paired with SA and CHO or PO. The relationship between synthesis factors and material properties relevant to degradation behaviour including *M̅_n_*, PDI, ester bonding, and T_g_ was measured through comprehensive statistical analyses to determine significant factors to properties outputs. Synthesis parameter variations (i.e. A:E ratio, M:C ratio, temperature, and time) were significantly correlated to all material properties, highlighting the opportunity for tunability in polymer properties. Material properties observed align with previously reported SA-based polyesters synthesized through ROCOP^19,24^. Comparison of PO to CHO copolymerization with SA under matched reaction conditions highlights the opportunity in this synthesis approach; the mean crude DP of PPS materials (DP = 24) was two times higher than that of PCS (DP = 13), yet PPS materials exhibited more rapid mass loss tied to notable differences in material properties.

The synthesis factor-response relationships, reduced according to statistical significance, enabled efficient appreciation for the relevance of specific input variables on output properties, the magnitude of influence that independent factors have on property variation, and interactive relationships between multiple factors. *M̅_n_*, PDI, and ester bonding, showed significant dependence on at least two synthesis factors. Notably, feed ratio factors A:E and M:C were consistently significant across material properties studied. At higher anhydride feed, chain propagation proceeded with more alternating insertions, yielding higher ester bonding and lower PDI. Higher epoxide feed increased the probability of double-epoxide insertion, which elevated *M̅_n_* but reduced structural fidelity through increased ether bond content undesirable to degradable polyester design^40,43^. Ester bonding trends diverged between PCS and PPS, reflecting differences in CHO and PO reactivity, with PCS showing more ether formation under limited initiation and PPS under non-limited initiation^39,43,44^. M:C ratio variation showed maxima *M̅_n_* behaviour, highlighting that achievement of high DP and sufficient initiation must be balanced when selecting catalyst concentration. Reaction temperature and time were notably less impactful on material properties, likely defined by the selected catalyst system operating in its effective range^14^. Nevertheless, it is noteworthy that time did play a significant interaction effect on multiple properties, indicative of the increased probability of transesterification and reduced alternating polymer structure with reaction progression^45,46^. The lack of plateau in PPS *M̅_n_* with the high (400:1) M:C ratio may suggest an opportunity for further optimization outside of the bounds investigated in this study^47^.

Variability in extraction of polyesters developed herein limited the predictive nature of regression models developed; the relatively low *M̅_n_* observed in all samples (i.e., <10kDa) led to variable extraction yields, where samples behaved both like polymers and like small molecules, leading to a bias of longer chain extraction as higher DP polymers are less soluble^48^. Future work could look to further optimize extraction for a material formulation of interest, including consideration of filtration approaches^40,49^ or extraction solvent iteration^29,43^. Nevertheless, the factor-response trends from crude materials were largely conserved in the extracted polyester properties, particularly for PDI and ester bonding. Notably, while not the primary focus of this study, the models for crude material were generally effective in reproducing *M̅_n_*, PDI, and alternating structure for a defined ideal case within the range of factors assessed. We expect the observed deviation in PPS PDI from predicted values can be attributed to deterioration of the PO reagent^50–53^, but this was not confirmed.

Degradation studies highlight the importance of monomer selection in the development of resorbable polyester material technologies. PPS formulations exhibited rapid degradation under accelerated conditions, where ∼50% of formulations degraded in 0.125 M NaOH, whereas PCS materials exhibited notably less mass loss in 1.0 M NaOH over the same time frame. This was reflected under neutral conditions; PPS materials of higher DP exhibited greater mass loss over 60 days. Interestingly, changes in *M̅_n_* and PDI in both formulations over this timeline were negligible, suggesting degradation proceeded through surface erosion^54,55^. The cyclohexyl ring of CHO likely impedes the rate of hydrolysis through steric hindrance and the electron density of the ring, as seen in comparative aliphatic and semi-aromatic degradation elsewhere^56^. Notably, our assessment suggests material properties (i.e., *M̅_n_*, PDI, alternating structure) provide opportunity to further tune resorption timelines, although additional assessment is needed to characterize hydrolysis of multiple materials at neutral conditions. The design opportunity in monomer selection through copolymerization has been well described elsewhere, including through condensation polymerization of di-acids and di-alcohols, as well as manipulation of ratios of lactide stereoisomers and ratios of lactide:glycolide in clinically applied poly(lactide-*co-*glycolide) polyester materials^4^. Indirect assessment of cytocompatibility presented no evidence of toxicity, providing the first benchmark to relevance of SA polyesters developed through ROCOP for biomaterial applications. Future work should extend the timelines of assessment and pursue study *in vivo* to appreciate other toxicity implications along with differences in resorption timelines driven by biologic microenvironmental factors.

## 5 CONCLUSIONS

This work demonstrates the systematic development of degradable polyesters via ROCOP of SA with CHO (PCS) or PO (PPS), guided by a CCD approach. Factor–response relationships revealed that feed ratios (A:E and M:C) consistently and significantly influenced *M̅_n_*, PDI, and alternating structure, while reaction time and temperature had comparatively minor effects. Importantly, PPS achieved higher degrees of polymerization than PCS but exhibited more rapid degradation under physiologic and accelerated conditions, highlighting how monomer selection directly tunes degradation outcomes. Both formulations demonstrated minimal cytotoxicity, supporting their promise for biomedical translation. The ROCOP approach described here establishes guidelines for rational design of resorbable biomaterials that combine the polymerization control of ring opening polymerization of cyclic lactones with the monomer diversity of copolymerization primarily described with polycondensation approaches. Future work could extend this methodology to other cyclic anhydrides (e.g., itaconic, citraconic) and epoxides to expand chemical diversity, and to alternative catalyst systems that may broaden control of polymer structure and properties. Collectively, these findings highlight the potential of ROCOP as a versatile synthesis strategy for next-generation degradable biomaterials with application-specific performance.

## 6 ASSOCIATED CONTENT

### Supporting Information

ANOVA tables for multi-variate regression models, equations for reduced regression models of extracted properties, representative NMR spectra and GPC traces used in analyses, summary of NMR analysis ranges, feature plots of extracted properties

## 7 AUTHOR INFORMATION

### Author Contributions

SM conceived the idea, completed the CCD and model validation experiments, and analyzed the results. KW & BW performed temporal degradation experiments and temporal molecular weight characterization and analysis. ZF performed cell toxicity experiments and analysis. AS & LDH supervised the entire project. All authors wrote, read and edited the manuscript and approved its final submission.

## 8 ACKNOWLEDGEMENTS

This research is supported by Natural Sciences and Engineering Research Council of Canada (NSERC) Discovery Grants (RGPIN-2021-02401; RGPIN-2022-03666), New Frontiers in Research Fund – Exploration Fund (NFRFE-2022-00313), and Canadian Institutes for Health Research Project Grant (PJT – 191692). We acknowledge infrastructure support from the Canadian Foundation for Innovation (JELF 42294, JELF 39824), Research Nova Scotia Research Opportunities Fund (2022-2400, 2020-1208), and NSERC Research Tools and Instrumentation Grant (RTI-2024-00128).

