## Supporting Information for "Systematic development of degradable polyester biomaterials via ring-opening copolymerization of succinic anhydride and epoxides"

Pages: 36

### 1 SUPPLEMENTAL EQUATIONS

PCS Extract $\overline{M}_n$ (g $mol^{-1}$) = 2553.46 - 366.56*A - 213.94*B + 370.28*D + 251.13*AB + 176.13*BD  Equation S1

PPS Extract $\overline{M}_n$ (g $mol^{-1}$) = 4804.66 - 566.72*A - 94.39*B + 426.33*D + 386.5*BD - 960.04*$B^2$  Equation S2

PCS Extract PDI = 1.40 - 0.1251*A - 0.0339*B + 0.0792*C + 0.0918*AB  Equation S3

PPS Extract PDI = 1.48 - 0.1795*A - 0.1022*B + 0.1049*AB  Equation S4

PCS Extract Ester Bond (%) = 93.14 + 2.87*A - 3.56*B + 2.07*D  Equation S5

PPS Extract Ester Bond (%) = 96.33 + 1.28*A + 2.77*B - 0.0606*C - 0.0406*D - 1.15*AB - 0.9987*CD  Equation S6

### 2 SUPPLEMENTAL FIGURES

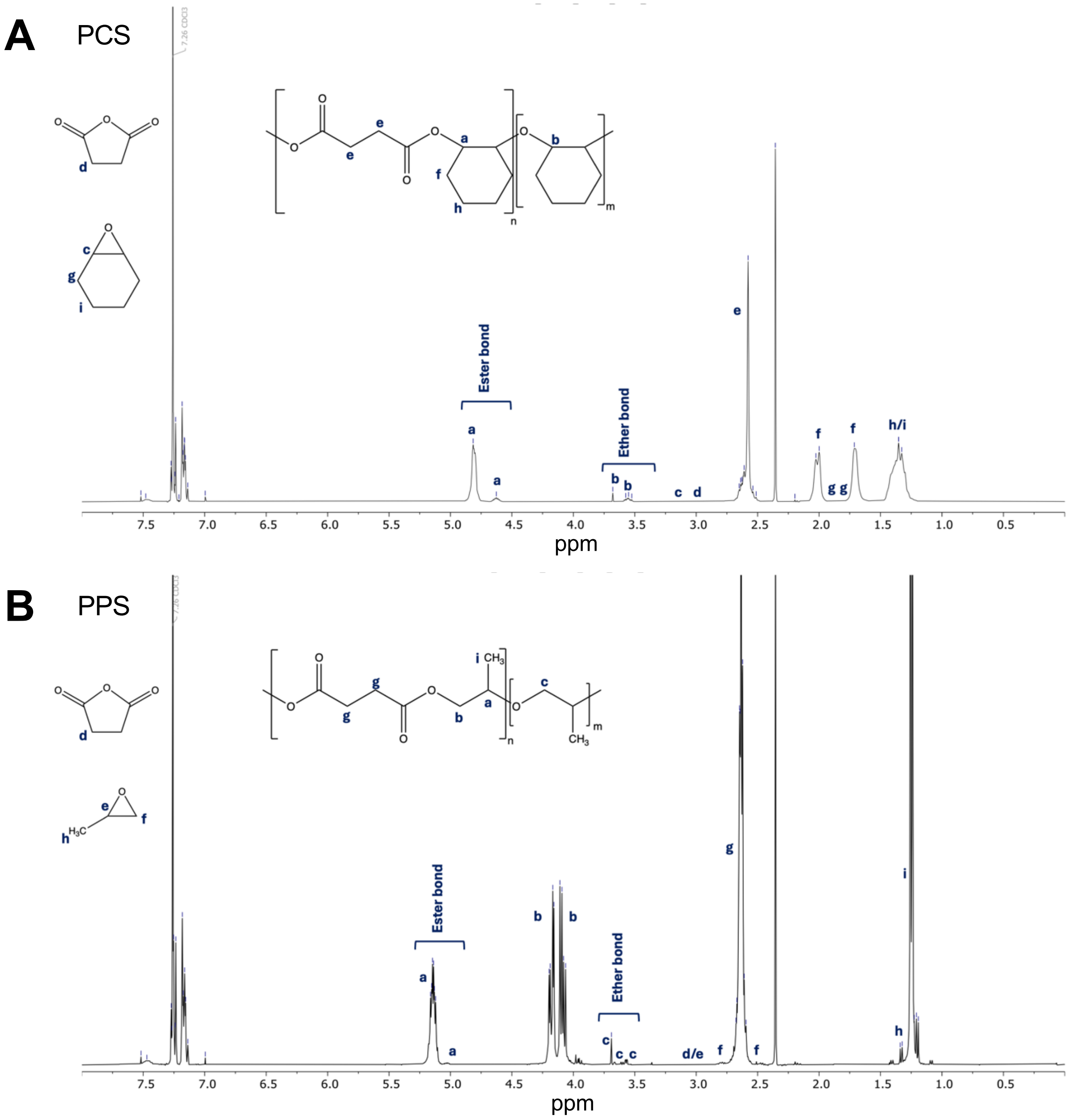


**Figure S1: Representative annotated $^1$H NMR spectra for (A) PCS and (B) PPS highlights peaks used for quantitative structural analyses**

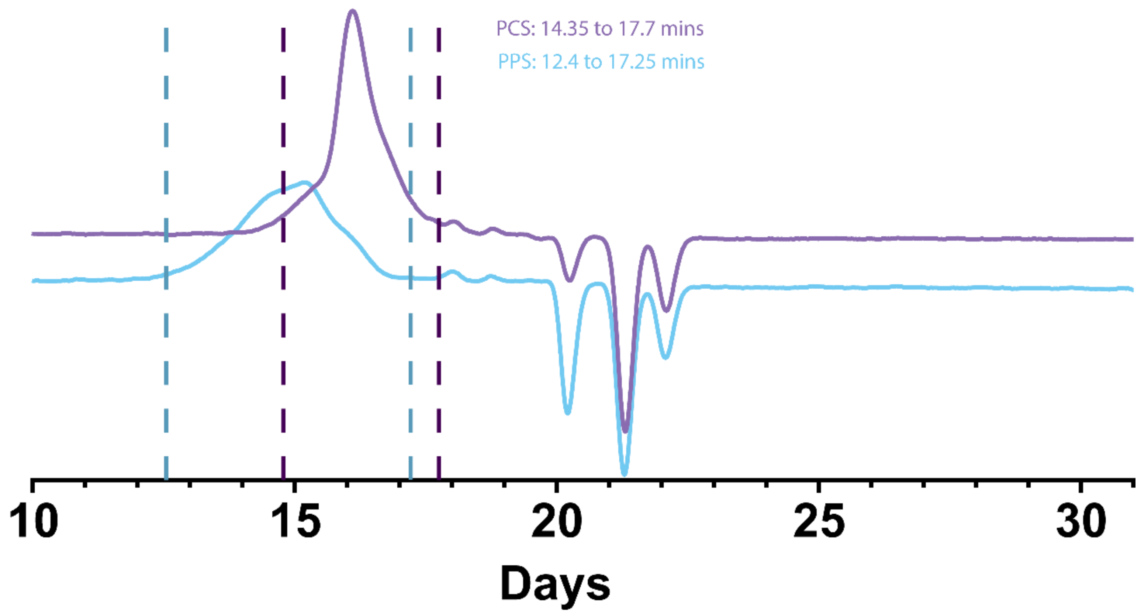


**Figure S2: Representative molecular weight trace demonstrating analysis approach for crude materials.**

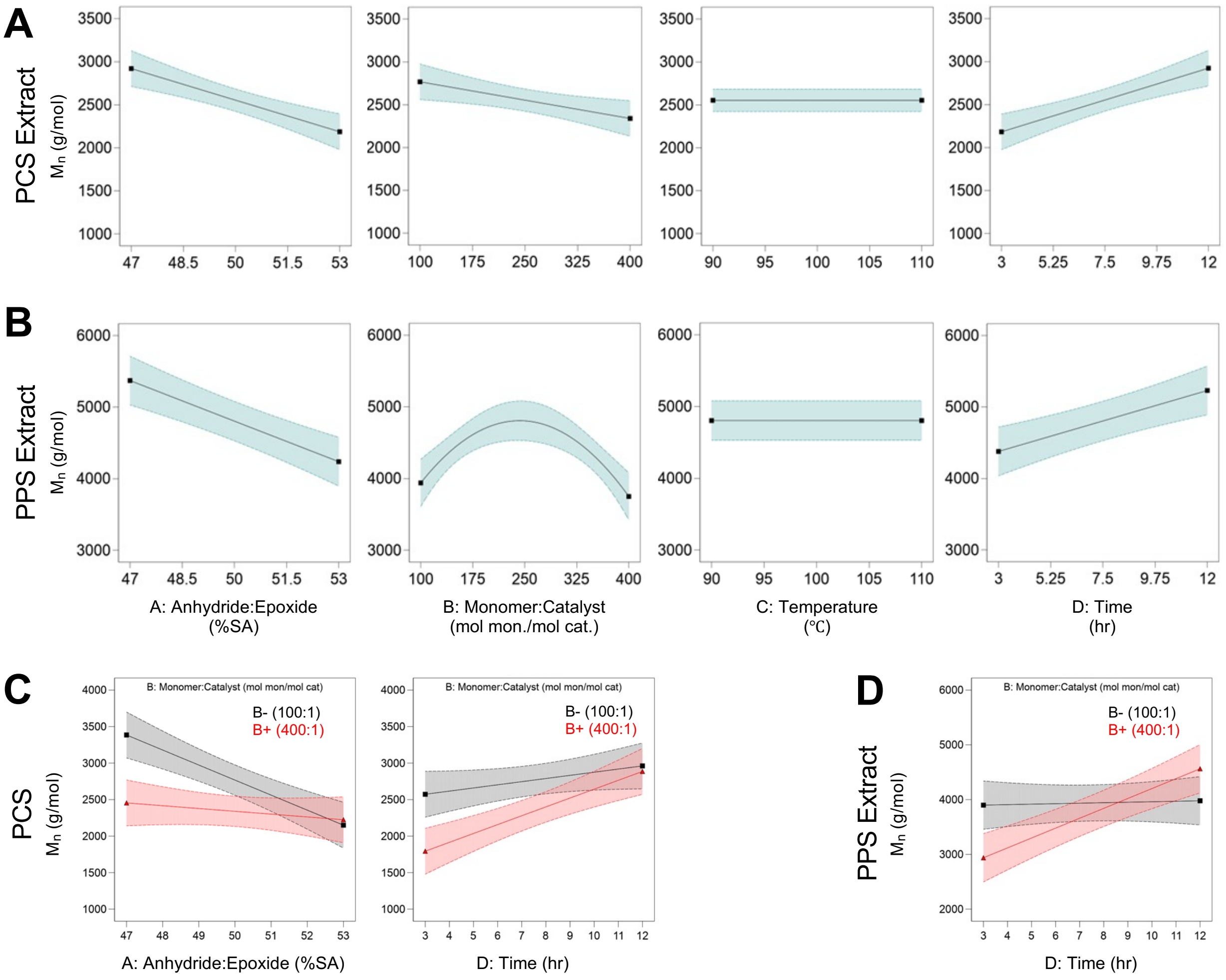


**Figure S3: Factor-response models highlight relationships between synthesis variables and extracted polyester $\bar{M}_n$.** (A-B) Visual representation of expected (A) PCS and (B) PPS extracted $\bar{M}_n$ as a function of synthesis factors. Data was empirically modelled and is presented within the high and low experimental bounds of each factor. Curves represent predictions from the reduced regression model shown across the high (+1) and low (-1) bounds of each factor while holding other factors at their midpoint (0). (C-D) Interaction plots illustrate the influence of significant interaction effects for (C) PCS and (D) PPS. Each plot shows two curves across the experimental range of the x-axis factor, with the interacting factor fixed either at its high (red) or low (grey) level. All other factors were held constant at their midpoint (coded value: 0). Shaded regions in all plots denote 95% confidence intervals.

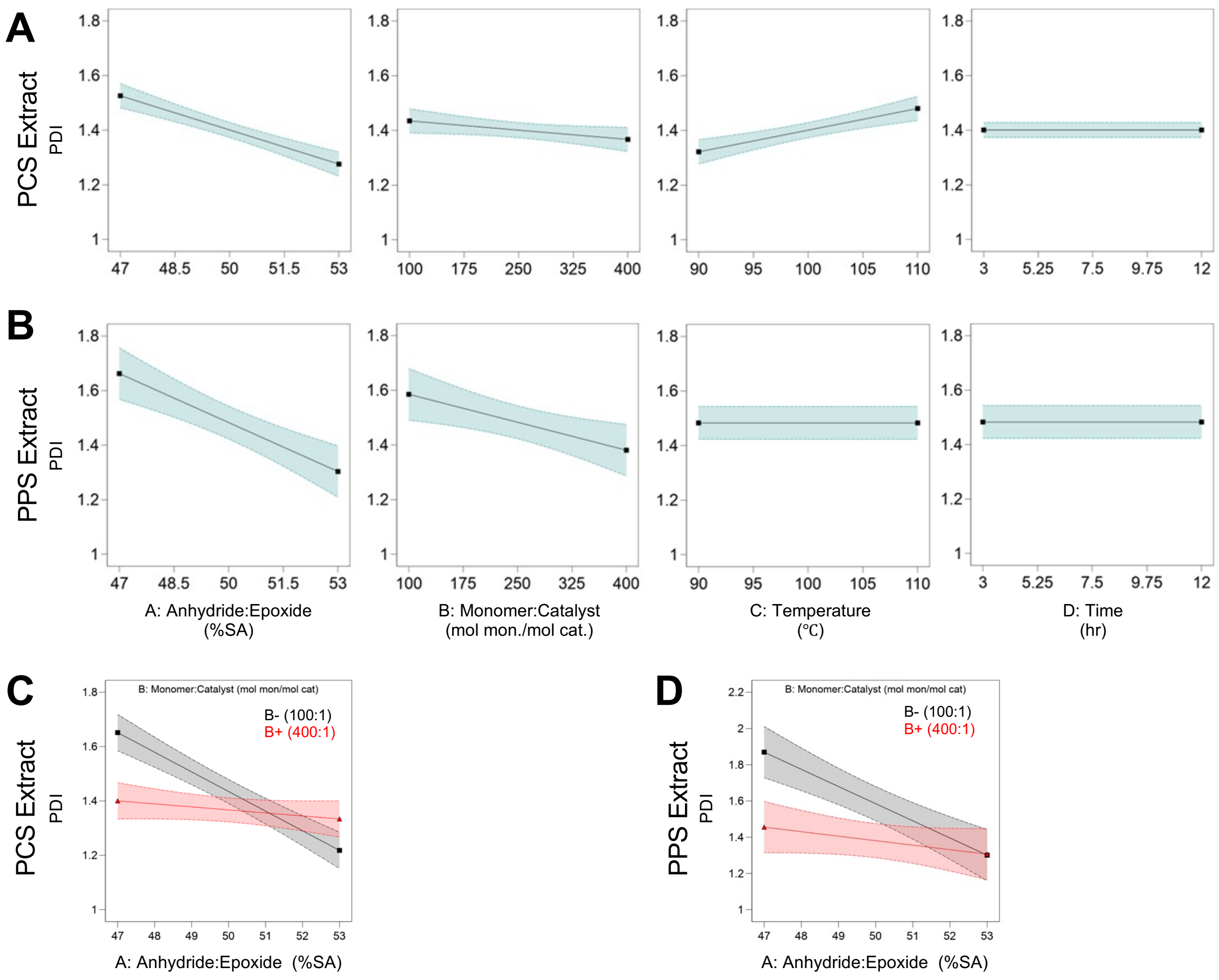


**Figure S4: Factor-response models highlight relationships between synthesis variables and extracted polyester PDI.** (A-B) Visual representation of expected (A) PCS and (B) PPS extracted PDI as a function of synthesis factors. Data was empirically modelled and is presented within the high and low experimental bounds of each factor. Curves represent predictions from the reduced regression model shown across the high (+1) and low (-1) bounds of each factor while holding other factors at their midpoint (0). (C-D) Interaction plots illustrate the influence of significant interaction effects for (C) PCS and (D) PPS. Each plot shows two curves across the experimental range of the x-axis factor, with the interacting factor fixed either at its high (red) or low (grey) level. All other factors were held constant at their midpoint (coded value: 0). Shaded regions in all plots denote 95% confidence intervals.

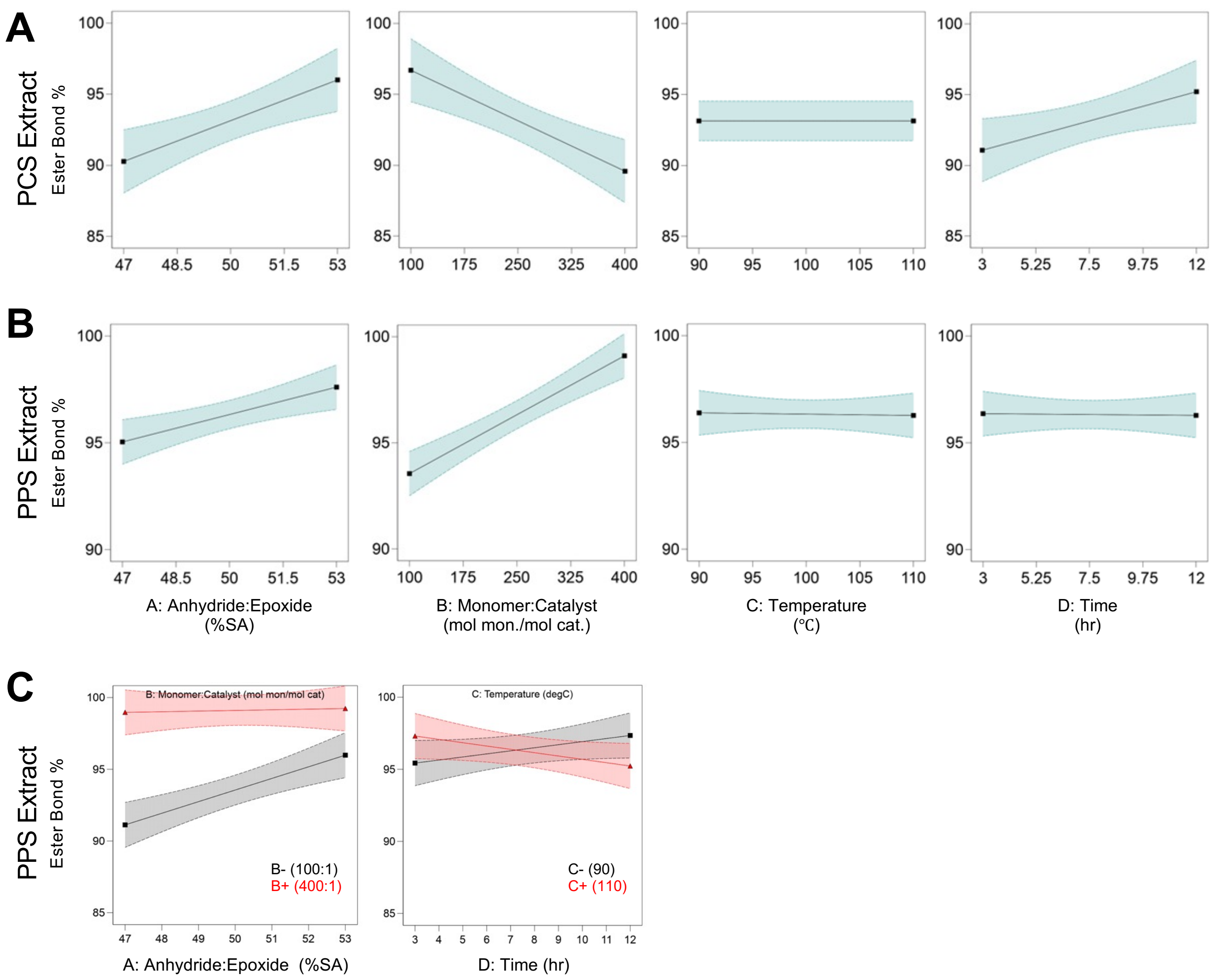


**Figure S5: Factor-response models highlight relationships between synthesis variables and extracted polyester alternating structure.** (A-B) Visual representation of expected (A) PCS and (B) PPS extracted ester bond frequency as a function of synthesis factors. Data was empirically modelled and is presented within the high and low experimental bounds of each factor. Curves represent predictions from the reduced regression model shown across the high (+1) and low (-1) bounds of each factor while holding other factors at their midpoint (0). (C-D) Interaction plots illustrate the influence of significant interaction effects for (C) PCS and (D) PPS. Each plot shows two curves across the experimental range of the x-axis factor, with the interacting factor fixed either at its high (red) or low (grey) level. All other factors were held constant at their midpoint (coded value: 0). Shaded regions in all plots denote 95% confidence intervals.

### 3 SUPPLEMENTAL TABLES

**Table S1: $^{1}$H-NMR integration regions used in quantification of alternating structure**

| PCS | | PPS | |
|---|---|---|---|
| **Assignment** | **Range (ppm)** | **Assignment** | **Range (ppm)** |
| Ester bond (a) | 4.90 .. 4.74 | Ester bond (a) | 5.26 .. 5.07 |
| Ester bond (a) | 4.68 .. 4.58 | Ester bond (a) | 5.06 .. 4.98 |
| Ether bond (b) | 3.70 .. 3.66 | b | 4.24 .. 4.03 |
| Ether bond (b) | 3.60 .. 3.51 | Ether bond (c) | 3.71 .. 3.68 |
| Ether bond (b) | 3.46 .. 3.30 | Ether bond (c) | 3.68 .. 3.65 |
| c | 3.14 .. 3.10 | Ether bond (c) | 3.62 .. 3.60 |
| d | 3.04 .. 2.96 | Ether bond (c) | 3.60 .. 3.59 |
| e | 2.69 .. 2.47 | Ether bond (c) | 3.59 .. 3.57 |
| f | 2.10 .. 1.96 | Ether bond (c) | 3.57 .. 3.55 |
| g | 1.95 .. 1.90 | Ether bond (c) | 3.54 .. 3.44 |
| g | 1.86 .. 1.77 | d/e | 3.05 .. 2.96 |
| f | 1.75 .. 1.63 | f | 2.85 .. 2.75 |
| h/i | 1.50 .. 1.14 | g | 2.75 .. 2.54 |
| | | f | 2.52 .. 2.43 |
| | | h | 1.35 .. 1.32 |
| | | i | 1.30 .. 1.22 |

**Table S2: PCS full model ANOVA of crude $\bar{M}_n$ response**

| Source | Sum of Squares | df | Mean Square | F-value | p-value |
|---|---|---|---|---|---|
| Model | 3.78E+06 | 14 | 2.70E+05 | 4.46 | 0.0053 |
| A-Anhydride:Epoxide | 2.06E+05 | 1 | 2.06E+05 | 3.41 | 0.0876 |
| B-Monomer:Catalyst | 2.42E+05 | 1 | 2.42E+05 | 4 | 0.0668 |
| C-Temperature | 34147.56 | 1 | 34147.56 | 0.5652 | 0.4656 |
| D-Time | 8.87E+05 | 1 | 8.87E+05 | 14.68 | 0.0021 |
| AB | 1.40E+05 | 1 | 1.40E+05 | 2.32 | 0.1513 |
| AC | 68251.56 | 1 | 68251.56 | 1.13 | 0.3072 |
| AD | 50737.56 | 1 | 50737.56 | 0.8398 | 0.3761 |
| BC | 1.80E+05 | 1 | 1.80E+05 | 2.98 | 0.108 |
| BD | 6.63E+05 | 1 | 6.63E+05 | 10.97 | 0.0056 |
| CD | 29670.06 | 1 | 29670.06 | 0.4911 | 0.4958 |
| $A^2$ | 2.94E+05 | 1 | 2.94E+05 | 4.87 | 0.046 |
| $B^2$ | 3.35E+05 | 1 | 3.35E+05 | 5.54 | 0.0349 |
| $C^2$ | 7143.97 | 1 | 7143.97 | 0.1182 | 0.7364 |
| $D^2$ | 3.51E+05 | 1 | 3.51E+05 | 5.81 | 0.0315 |
| Residual | 7.85E+05 | 13 | 60416.06 | | |
| Lack of Fit | 6.56E+05 | 10 | 65579.78 | 1.52 | 0.4042 |
| Pure Error | 1.30E+05 | 3 | 43203.67 | | |
| Cor Total | 4.56E+06 | 27 | | | |

**Table S3: PCS reduced model ANOVA of crude $\bar{M}_n$ response**

| Source | Sum of Squares | df | Mean Square | F-value | p-value |
|---|---|---|---|---|---|
| Model | 3.27E+06 | 7 | 4.66E+05 | 7.2 | 0.0002 |
| A-Anhydride:Epoxide | 2.06E+05 | 1 | 2.06E+05 | 3.18 | 0.0897 |
| B-Monomer:Catalyst | 2.42E+05 | 1 | 2.42E+05 | 3.73 | 0.0677 |
| D-Time | 8.87E+05 | 1 | 8.87E+05 | 13.69 | 0.0014 |
| BD | 6.63E+05 | 1 | 6.63E+05 | 10.23 | 0.0045 |
| $A^2$ | 3.51E+05 | 1 | 3.51E+05 | 5.41 | 0.0306 |
| $B^2$ | 3.35E+05 | 1 | 3.35E+05 | 5.18 | 0.034 |
| $D^2$ | 3.52E+05 | 1 | 3.52E+05 | 5.43 | 0.0303 |
| Residual | 1.30E+06 | 20 | 64789.26 | | |
| Lack of Fit | 1.17E+06 | 17 | 68598.48 | 1.59 | 0.3943 |
| Pure Error | 1.30E+05 | 3 | 43203.67 | | |
| Cor Total | 4.56E+06 | 27 | | | |

**Table S4: PCS full model ANOVA of extract $\bar{M}_n$ response**

| Source | Sum of Squares | df | Mean Square | F-value | p-value |
|---|---|---|---|---|---|
| Model | 8.48E+06 | 14 | 6.06E+05 | 6.9 | 0.0006 |
| A-Anhydride:Epoxide | 2.42E+06 | 1 | 2.42E+06 | 27.54 | 0.0002 |
| B-Monomer:Catalyst | 8.24E+05 | 1 | 8.24E+05 | 9.38 | 0.0091 |
| C-Temperature | 13068.06 | 1 | 13068.06 | 0.1488 | 0.7059 |
| D-Time | 2.47E+06 | 1 | 2.47E+06 | 28.1 | 0.0001 |
| AB | 1.01E+06 | 1 | 1.01E+06 | 11.49 | 0.0048 |
| AC | 50400.25 | 1 | 50400.25 | 0.5739 | 0.4622 |
| AD | 16770.25 | 1 | 16770.25 | 0.1909 | 0.6693 |
| BC | 3.69E+05 | 1 | 3.69E+05 | 4.2 | 0.0611 |
| BD | 4.96E+05 | 1 | 4.96E+05 | 5.65 | 0.0335 |
| CD | 3.62E+05 | 1 | 3.62E+05 | 4.12 | 0.0634 |
| $A^2$ | 58960.65 | 1 | 58960.65 | 0.6713 | 0.4274 |
| $B^2$ | 1.12E+05 | 1 | 1.12E+05 | 1.27 | 0.2793 |
| $C^2$ | 1785.75 | 1 | 1785.75 | 0.0203 | 0.8888 |
| $D^2$ | 68795.39 | 1 | 68795.39 | 0.7833 | 0.3922 |
| Residual | 1.14E+06 | 13 | 87827.62 | | |
| Lack of Fit | 4.56E+05 | 10 | 45578 | 0.1993 | 0.9776 |
| Pure Error | 6.86E+05 | 3 | 2.29E+05 | | |
| Cor Total | 9.63E+06 | 27 | | | |

**Table S5: PCS reduced model ANOVA of extract $\bar{M}_n$ response**

| Source | Sum of Squares | df | Mean Square | F-value | p-value |
|---|---|---|---|---|---|
| Model | 7.22E+06 | 5 | 1.44E+06 | 13.18 | < 0.0001 |
| A-Anhydride:Epoxide | 2.42E+06 | 1 | 2.42E+06 | 22.08 | 0.0001 |
| B-Monomer:Catalyst | 8.24E+05 | 1 | 8.24E+05 | 7.52 | 0.0119 |
| D-Time | 2.47E+06 | 1 | 2.47E+06 | 22.53 | < 0.0001 |
| AB | 1.01E+06 | 1 | 1.01E+06 | 9.21 | 0.0061 |
| BD | 4.96E+05 | 1 | 4.96E+05 | 4.53 | 0.0447 |
| Residual | 2.41E+06 | 22 | 1.10E+05 | | |
| Lack of Fit | 1.72E+06 | 19 | 90722.87 | 0.3968 | 0.9113 |
| Pure Error | 6.86E+05 | 3 | 2.29E+05 | | |
| Cor Total | 9.63E+06 | 27 | | | |
| Model | 7.22E+06 | 5 | 1.44E+06 | 13.18 | < 0.0001 |

**Table S6: PPS full model ANOVA of crude $\bar{M}_n$ response**

| Source | Sum of Squares | df | Mean Square | F-value | p-value |
|---|---|---|---|---|---|
| Model | 1.48E+07 | 14 | 1.06E+06 | 5.8 | 0.0015 |
| A-Anhydride:Epoxide | 4.16E+06 | 1 | 4.16E+06 | 22.85 | 0.0004 |
| B-Monomer:Catalyst | 0.5 | 1 | 0.5 | 2.75E-06 | 0.9987 |
| C-Temperature | 1.39E+05 | 1 | 1.39E+05 | 0.7632 | 0.3982 |
| D-Time | 3.73E+06 | 1 | 3.73E+06 | 20.48 | 0.0006 |
| AB | 49284 | 1 | 49284 | 0.2709 | 0.6115 |
| AC | 24964 | 1 | 24964 | 0.1372 | 0.717 |
| AD | 8.05E+05 | 1 | 8.05E+05 | 4.42 | 0.0555 |
| BC | 88506.25 | 1 | 88506.25 | 0.4864 | 0.4978 |
| BD | 2.79E+06 | 1 | 2.79E+06 | 15.32 | 0.0018 |
| CD | 6162.25 | 1 | 6162.25 | 0.0339 | 0.8568 |
| $A^2$ | 54045.86 | 1 | 54045.86 | 0.297 | 0.595 |
| $B^2$ | 1.59E+06 | 1 | 1.59E+06 | 8.74 | 0.0111 |
| $C^2$ | 3.55 | 1 | 3.55 | 0 | 0.9965 |
| $D^2$ | 1.16E+05 | 1 | 1.16E+05 | 0.6381 | 0.4387 |
| Residual | 2.37E+06 | 13 | 1.82E+05 | | |
| Lack of Fit | 1.63E+06 | 10 | 1.63E+05 | 0.6632 | 0.7281 |
| Pure Error | 7.37E+05 | 3 | 2.46E+05 | | |
| Cor Total | 2.42E+07 | 28 | | | |

**Table S7: PPS reduced model ANOVA of crude $\bar{M}_n$ response**

| Source | Sum of Squares | df | Mean Square | F-value | p-value |
|---|---|---|---|---|---|
| Model | 1.43E+07 | 6 | 2.39E+06 | 17.83 | < 0.0001 |
| A-Anhydride:Epoxide | 4.16E+06 | 1 | 4.16E+06 | 31.03 | < 0.0001 |
| B-Monomer:Catalyst | 0.5 | 1 | 0.5 | 3.73E-06 | 0.9985 |
| D-Time | 3.73E+06 | 1 | 3.73E+06 | 27.82 | < 0.0001 |
| AD | 8.05E+05 | 1 | 8.05E+05 | 6 | 0.0231 |
| BD | 2.79E+06 | 1 | 2.79E+06 | 20.8 | 0.0002 |
| $B^2$ | 2.86E+06 | 1 | 2.86E+06 | 21.32 | 0.0001 |
| Residual | 2.81E+06 | 21 | 1.34E+05 | | |
| Lack of Fit | 2.08E+06 | 18 | 1.15E+05 | 0.4699 | 0.8677 |
| Pure Error | 7.37E+05 | 3 | 2.46E+05 | | |
| Cor Total | 2.42E+07 | 28 | | | |

**Table S8: PPS full model ANOVA of extract $\bar{M}_n$ response**

| Source | Sum of Squares | df | Mean Square | F-value | p-value |
|---|---|---|---|---|---|
| Model | 1.70E+07 | 14 | 1.21E+06 | 7.24 | 0.0005 |
| A-Anhydride:Epoxide | 5.78E+06 | 1 | 5.78E+06 | 34.55 | < 0.0001 |
| B-Monomer:Catalyst | 1.60E+05 | 1 | 1.60E+05 | 0.9584 | 0.3455 |
| C-Temperature | 94322.72 | 1 | 94322.72 | 0.5637 | 0.4662 |
| D-Time | 3.27E+06 | 1 | 3.27E+06 | 19.55 | 0.0007 |
| AB | 4.82E+05 | 1 | 4.82E+05 | 2.88 | 0.1133 |
| AC | 2.10E+05 | 1 | 2.10E+05 | 1.25 | 0.2831 |
| AD | 4.85E+05 | 1 | 4.85E+05 | 2.9 | 0.1124 |
| BC | 1.44E+05 | 1 | 1.44E+05 | 0.8607 | 0.3705 |
| BD | 2.39E+06 | 1 | 2.39E+06 | 14.28 | 0.0023 |
| CD | 240.25 | 1 | 240.25 | 0.0014 | 0.9704 |
| $A^2$ | 15819.12 | 1 | 15819.12 | 0.0945 | 0.7634 |
| $B^2$ | 1.94E+06 | 1 | 1.94E+06 | 11.62 | 0.0047 |
| $C^2$ | 608.14 | 1 | 608.14 | 0.0036 | 0.9528 |
| $D^2$ | 1.19E+05 | 1 | 1.19E+05 | 0.7126 | 0.4138 |
| Residual | 2.18E+06 | 13 | 1.67E+05 | | |
| Lack of Fit | 1.38E+06 | 10 | 1.38E+05 | 0.5205 | 0.8098 |
| Pure Error | 7.95E+05 | 3 | 2.65E+05 | | |
| Cor Total | 2.24E+07 | 28 | | | |

**Table S9: PPS reduced model ANOVA of extract $\bar{M}_n$ (g mol$^{-1}$) response**

| Source | Sum of Squares | df | Mean Square | F-value | p-value |
|---|---|---|---|---|---|
| Source | Sum of Squares | df | Mean Square | F-value | p-value |
| Block | 3.24E+06 | 1 | 3.24E+06 | | |
| Model | 1.54E+07 | 5 | 3.08E+06 | 18.2 | < 0.0001 |
| A-Anhydride:Epoxide | 5.78E+06 | 1 | 5.78E+06 | 34.15 | < 0.0001 |
| B-Monomer:Catalyst | 1.60E+05 | 1 | 1.60E+05 | 0.9474 | 0.341 |
| D-Time | 3.27E+06 | 1 | 3.27E+06 | 19.33 | 0.0002 |
| BD | 2.39E+06 | 1 | 2.39E+06 | 14.12 | 0.0011 |
| $B^2$ | 3.80E+06 | 1 | 3.80E+06 | 22.47 | < 0.0001 |
| Residual | 3.72E+06 | 22 | 1.69E+05 | | |
| Lack of Fit | 2.93E+06 | 19 | 1.54E+05 | 0.5814 | 0.8032 |
| Pure Error | 7.95E+05 | 3 | 2.65E+05 | | |
| Cor Total | 2.24E+07 | 28 | | | |

**Table S10: PCS full model ANOVA of crude PDI response**

| Source | Sum of Squares | df | Mean Square | F-value | p-value |
|---|---|---|---|---|---|
| Model | 0.65 | 14 | 0.0464 | 13.59 | < 0.0001 |
| A-Anhydride:Epoxide | 0.2699 | 1 | 0.2699 | 79.02 | < 0.0001 |
| B-Monomer:Catalyst | 0.0867 | 1 | 0.0867 | 25.38 | 0.0002 |
| C-Temperature | 0.0175 | 1 | 0.0175 | 5.12 | 0.0414 |
| D-Time | 0.0698 | 1 | 0.0698 | 20.44 | 0.0006 |
| AB | 0.1223 | 1 | 0.1223 | 35.82 | < 0.0001 |
| AC | 0.007 | 1 | 0.007 | 2.05 | 0.1754 |
| AD | 0.0358 | 1 | 0.0358 | 10.49 | 0.0065 |
| BC | 0.0032 | 1 | 0.0032 | 0.9265 | 0.3533 |
| BD | 0.0027 | 1 | 0.0027 | 0.7994 | 0.3875 |
| CD | 0 | 1 | 0 | 0.0114 | 0.9165 |
| $A^2$ | 0.0097 | 1 | 0.0097 | 2.84 | 0.116 |
| $B^2$ | 0 | 1 | 0 | 0.0058 | 0.9402 |
| $C^2$ | 0.0016 | 1 | 0.0016 | 0.4639 | 0.5078 |
| $D^2$ | 0.0004 | 1 | 0.0004 | 0.1127 | 0.7424 |
| Residual | 0.0444 | 13 | 0.0034 | | |
| Lack of Fit | 0.044 | 10 | 0.0044 | 35.24 | 0.0069 |
| Pure Error | 0.0004 | 3 | 0.0001 | | |
| Cor Total | 0.6944 | 27 | | | |

**Table S11: PCS reduced model ANOVA of crude PDI response**

| Source | Sum of Squares | df | Mean Square | F-value | p-value |
|---|---|---|---|---|---|
| Model | 0.602 | 6 | 0.1003 | 22.8 | < 0.0001 |
| A-Anhydride:Epoxide | 0.2699 | 1 | 0.2699 | 61.33 | < 0.0001 |
| B-Monomer:Catalyst | 0.0867 | 1 | 0.0867 | 19.7 | 0.0002 |
| C-Temperature | 0.0175 | 1 | 0.0175 | 3.97 | 0.0594 |
| D-Time | 0.0698 | 1 | 0.0698 | 15.87 | 0.0007 |
| AB | 0.1223 | 1 | 0.1223 | 27.8 | < 0.0001 |
| AD | 0.0358 | 1 | 0.0358 | 8.14 | 0.0095 |
| Residual | 0.0924 | 21 | 0.0044 | | |
| Lack of Fit | 0.092 | 18 | 0.0051 | 40.93 | 0.0054 |
| Pure Error | 0.0004 | 3 | 0.0001 | | |
| Cor Total | 0.6944 | 27 | | | |

**Table S12: PCS full model ANOVA of extract PDI response**

| Source | Sum of Squares | df | Mean Square | F-value | p-value |
|---|---|---|---|---|---|
| Model | 0.6108 | 14 | 0.0436 | 10.49 | < 0.0001 |
| A-Anhydride:Epoxide | 0.2815 | 1 | 0.2815 | 67.7 | < 0.0001 |
| B-Monomer:Catalyst | 0.0207 | 1 | 0.0207 | 4.97 | 0.044 |
| C-Temperature | 0.113 | 1 | 0.113 | 27.17 | 0.0002 |
| D-Time | 0.0044 | 1 | 0.0044 | 1.05 | 0.3247 |
| AB | 0.1347 | 1 | 0.1347 | 32.39 | < 0.0001 |
| AC | 0.0016 | 1 | 0.0016 | 0.3753 | 0.5507 |
| AD | 0.0044 | 1 | 0.0044 | 1.05 | 0.3247 |
| BC | 0.0086 | 1 | 0.0086 | 2.08 | 0.1729 |
| BD | 0.0029 | 1 | 0.0029 | 0.6884 | 0.4217 |
| CD | 0.0069 | 1 | 0.0069 | 1.66 | 0.2205 |
| $A^2$ | 0.0073 | 1 | 0.0073 | 1.76 | 0.2072 |
| $B^2$ | 0.0016 | 1 | 0.0016 | 0.3812 | 0.5476 |
| $C^2$ | 0 | 1 | 0 | 0.003 | 0.9569 |
| $D^2$ | 1.14E-07 | 1 | 1.14E-07 | 0 | 0.9959 |
| Residual | 0.0541 | 13 | 0.0042 | | |
| Lack of Fit | 0.0512 | 10 | 0.0051 | 5.46 | 0.0945 |
| Pure Error | 0.0028 | 3 | 0.0009 | | |
| Cor Total | 0.6648 | 27 | | | |

**Table S13: PCS reduced model ANOVA of extract PDI response**

| Source | Sum of Squares | df | Mean Square | F-value | p-value |
|---|---|---|---|---|---|
| Model | 0.5498 | 4 | 0.1375 | 27.49 | < 0.0001 |
| A-Anhydride:Epoxide | 0.2815 | 1 | 0.2815 | 56.3 | < 0.0001 |
| B-Monomer:Catalyst | 0.0207 | 1 | 0.0207 | 4.13 | 0.0537 |
| C-Temperature | 0.113 | 1 | 0.113 | 22.6 | < 0.0001 |
| AB | 0.1347 | 1 | 0.1347 | 26.94 | < 0.0001 |
| Residual | 0.115 | 23 | 0.005 | | |
| Lack of Fit | 0.1122 | 20 | 0.0056 | 5.98 | 0.0828 |
| Pure Error | 0.0028 | 3 | 0.0009 | | |
| Cor Total | 0.6648 | 27 | | | |

**Table S14: PPS full model ANOVA of crude PDI response**

| Source | Sum of Squares | df | Mean Square | F-value | p-value |
|---|---|---|---|---|---|
| Model | 1.6 | 14 | 0.1144 | 7.81 | 0.0003 |
| A-Anhydride:Epoxide | 0.7462 | 1 | 0.7462 | 50.92 | < 0.0001 |
| B-Monomer:Catalyst | 0.2913 | 1 | 0.2913 | 19.88 | 0.0006 |
| C-Temperature | 0.0821 | 1 | 0.0821 | 5.61 | 0.0341 |
| D-Time | 0.0202 | 1 | 0.0202 | 1.38 | 0.2614 |
| AB | 0.328 | 1 | 0.328 | 22.39 | 0.0004 |
| AC | 0.0244 | 1 | 0.0244 | 1.67 | 0.2193 |
| AD | 0.0016 | 1 | 0.0016 | 0.1078 | 0.7479 |
| BC | 0.0019 | 1 | 0.0019 | 0.1306 | 0.7236 |
| BD | 0.028 | 1 | 0.028 | 1.91 | 0.1904 |
| CD | 0.0099 | 1 | 0.0099 | 0.6722 | 0.4271 |
| $A^2$ | 0.0658 | 1 | 0.0658 | 4.49 | 0.0539 |
| $B^2$ | 0.0101 | 1 | 0.0101 | 0.69 | 0.4212 |
| $C^2$ | 0.0004 | 1 | 0.0004 | 0.0267 | 0.8726 |
| $D^2$ | 0.0058 | 1 | 0.0058 | 0.3943 | 0.5409 |
| Residual | 0.1905 | 13 | 0.0147 | | |
| Lack of Fit | 0.1038 | 10 | 0.0104 | 0.3589 | 0.9043 |
| Pure Error | 0.0867 | 3 | 0.0289 | | |
| Cor Total | 1.79 | 28 | | | |

**Table S15: PPS reduced model ANOVA of crude PDI response**

| Source | Sum of Squares | df | Mean Square | F-value | p-value |
|---|---|---|---|---|---|
| Model | 1.45 | 4 | 0.3619 | 24.13 | < 0.0001 |
| A-Anhydride:Epoxide | 0.7462 | 1 | 0.7462 | 49.75 | < 0.0001 |
| B-Monomer:Catalyst | 0.2913 | 1 | 0.2913 | 19.42 | 0.0002 |
| C-Temperature | 0.0821 | 1 | 0.0821 | 5.48 | 0.0283 |
| AB | 0.328 | 1 | 0.328 | 21.87 | 0.0001 |
| Residual | 0.345 | 23 | 0.015 | | |
| Lack of Fit | 0.2582 | 20 | 0.0129 | 0.4466 | 0.8849 |
| Pure Error | 0.0867 | 3 | 0.0289 | | |
| Cor Total | 1.79 | 28 | | | |

**Table S16: PPS full model ANOVA of extract PDI response**

| Source | Sum of Squares | df | Mean Square | F-value | p-value |
|---|---|---|---|---|---|
| Model | 1.19 | 14 | 0.0853 | 3.9 | 0.0095 |
| A-Anhydride:Epoxide | 0.58 | 1 | 0.58 | 26.53 | 0.0002 |
| B-Monomer:Catalyst | 0.1879 | 1 | 0.1879 | 8.59 | 0.0117 |
| C-Temperature | 0.0524 | 1 | 0.0524 | 2.4 | 0.1456 |
| D-Time | 0.0506 | 1 | 0.0506 | 2.31 | 0.1523 |
| AB | 0.1762 | 1 | 0.1762 | 8.06 | 0.014 |
| AC | 0.0077 | 1 | 0.0077 | 0.3522 | 0.563 |
| AD | 0.0081 | 1 | 0.0081 | 0.3726 | 0.5521 |
| BC | 0.0066 | 1 | 0.0066 | 0.302 | 0.592 |
| BD | 0.0423 | 1 | 0.0423 | 1.94 | 0.1874 |
| CD | 0.0002 | 1 | 0.0002 | 0.0069 | 0.9352 |
| $A^2$ | 0.0526 | 1 | 0.0526 | 2.41 | 0.1448 |
| $B^2$ | 0.0157 | 1 | 0.0157 | 0.7184 | 0.412 |
| $C^2$ | 0.0079 | 1 | 0.0079 | 0.3615 | 0.558 |
| $D^2$ | 0.0151 | 1 | 0.0151 | 0.6914 | 0.4207 |
| Residual | 0.2842 | 13 | 0.0219 | | |
| Lack of Fit | 0.1977 | 10 | 0.0198 | 0.6862 | 0.7156 |
| Pure Error | 0.0865 | 3 | 0.0288 | | |
| Cor Total | 1.5 | 28 | | | |

**Table S17: PPS reduced model ANOVA of extract PDI response**

| Source | Sum of Squares | df | Mean Square | F-value | p-value |
|---|---|---|---|---|---|
| Model | 0.944 | 3 | 0.3147 | 14.12 | < 0.0001 |
| A-Anhydride:Epoxide | 0.58 | 1 | 0.58 | 26.03 | < 0.0001 |
| B-Monomer:Catalyst | 0.1879 | 1 | 0.1879 | 8.43 | 0.0078 |
| AB | 0.1762 | 1 | 0.1762 | 7.91 | 0.0097 |
| Residual | 0.5348 | 24 | 0.0223 | | |
| Lack of Fit | 0.4483 | 21 | 0.0213 | 0.7408 | 0.7147 |
| Pure Error | 0.0865 | 3 | 0.0288 | | |
| Cor Total | 1.5 | 28 | | | |

**Table S18: PCS full model ANOVA of crude ester bond response**

| Source | Sum of Squares | df | Mean Square | F-value | p-value |
|---|---|---|---|---|---|
| Model | 595.35 | 14 | 42.52 | 10.69 | < 0.0001 |
| A-Anhydride:Epoxide | 133.12 | 1 | 133.12 | 33.48 | < 0.0001 |
| B-Monomer:Catalyst | 252.98 | 1 | 252.98 | 63.62 | < 0.0001 |
| C-Temperature | 2.61 | 1 | 2.61 | 0.6575 | 0.432 |
| D-Time | 74.91 | 1 | 74.91 | 18.84 | 0.0008 |
| AB | 19.51 | 1 | 19.51 | 4.91 | 0.0452 |
| AC | 3.41 | 1 | 3.41 | 0.8584 | 0.3711 |
| AD | 11.54 | 1 | 11.54 | 2.9 | 0.1122 |
| BC | 2.23 | 1 | 2.23 | 0.5602 | 0.4675 |
| BD | 44.12 | 1 | 44.12 | 11.1 | 0.0054 |
| CD | 5.19 | 1 | 5.19 | 1.3 | 0.274 |
| $A^2$ | 0.3706 | 1 | 0.3706 | 0.0932 | 0.765 |
| $B^2$ | 5.32 | 1 | 5.32 | 1.34 | 0.2683 |
| $C^2$ | 1.28 | 1 | 1.28 | 0.3215 | 0.5804 |
| $D^2$ | 13.13 | 1 | 13.13 | 3.3 | 0.0923 |
| Residual | 51.69 | 13 | 3.98 | | |
| Lack of Fit | 50.74 | 10 | 5.07 | 16.01 | 0.0215 |
| Pure Error | 0.9507 | 3 | 0.3169 | | |
| Cor Total | 647.04 | 27 | | | |

**Table S19: PCS reduced model ANOVA of crude ester bond response**

| Source | Sum of Squares | df | Mean Square | F-value | p-value |
|---|---|---|---|---|---|
| Model | 524.64 | 5 | 104.93 | 18.86 | < 0.0001 |
| A-Anhydride:Epoxide | 133.12 | 1 | 133.12 | 23.93 | < 0.0001 |
| B-Monomer:Catalyst | 252.98 | 1 | 252.98 | 45.47 | < 0.0001 |
| D-Time | 74.91 | 1 | 74.91 | 13.46 | 0.0013 |
| AB | 19.51 | 1 | 19.51 | 3.51 | 0.0744 |
| BD | 44.12 | 1 | 44.12 | 7.93 | 0.0101 |
| Residual | 122.4 | 22 | 5.56 | | |
| Lack of Fit | 121.45 | 19 | 6.39 | 20.17 | 0.0151 |
| Pure Error | 0.9507 | 3 | 0.3169 | | |
| Cor Total | 647.04 | 27 | | | |

**Table S20: PCS full model ANOVA of extract ester bond response**

| Source | Sum of Squares | df | Mean Square | F-value | p-value |
|---|---|---|---|---|---|
| Model | 575.14 | 14 | 41.08 | 2.92 | 0.0305 |
| A-Anhydride:Epoxide | 147.92 | 1 | 147.92 | 10.53 | 0.0064 |
| B-Monomer:Catalyst | 228.12 | 1 | 228.12 | 16.24 | 0.0014 |
| C-Temperature | 0.6013 | 1 | 0.6013 | 0.0428 | 0.8393 |
| D-Time | 77.05 | 1 | 77.05 | 5.48 | 0.0358 |
| AB | 1.6 | 1 | 1.6 | 0.1139 | 0.7411 |
| AC | 11.66 | 1 | 11.66 | 0.8302 | 0.3788 |
| AD | 16.2 | 1 | 16.2 | 1.15 | 0.3024 |
| BC | 8.97 | 1 | 8.97 | 0.6386 | 0.4386 |
| BD | 44.29 | 1 | 44.29 | 3.15 | 0.0992 |
| CD | 15.33 | 1 | 15.33 | 1.09 | 0.3152 |
| $A^2$ | 0.0058 | 1 | 0.0058 | 0.0004 | 0.9841 |
| $B^2$ | 10.6 | 1 | 10.6 | 0.7548 | 0.4007 |
| $C^2$ | 0.3116 | 1 | 0.3116 | 0.0222 | 0.8839 |
| $D^2$ | 0.0298 | 1 | 0.0298 | 0.0021 | 0.964 |
| Residual | 182.61 | 13 | 14.05 | | |
| Lack of Fit | 180.87 | 10 | 18.09 | 31.34 | 0.0081 |
| Pure Error | 1.73 | 3 | 0.5772 | | |
| Cor Total | 757.74 | 27 | | | |

**Table S21: PCS reduced model ANOVA of extract ester bond response**

| Source | Sum of Squares | df | Mean Square | F-value | p-value |
|---|---|---|---|---|---|
| Model | 453.09 | 3 | 151.03 | 11.9 | < 0.0001 |
| A-Anhydride:Epoxide | 147.92 | 1 | 147.92 | 11.65 | 0.0023 |
| B-Monomer:Catalyst | 228.12 | 1 | 228.12 | 17.97 | 0.0003 |
| D-Time | 77.05 | 1 | 77.05 | 6.07 | 0.0213 |
| Residual | 304.65 | 24 | 12.69 | | |
| Lack of Fit | 302.92 | 21 | 14.42 | 24.99 | 0.011 |
| Pure Error | 1.73 | 3 | 0.5772 | | |
| Cor Total | 757.74 | 27 | | | |

**Table S22: PPS full model ANOVA of crude ester bond response**

| Source | Sum of Squares | df | Mean Square | F-value | p-value |
|---|---|---|---|---|---|
| Model | 257.8 | 14 | 18.41 | 6.59 | 0.0008 |
| A-Anhydride:Epoxide | 45.09 | 1 | 45.09 | 16.14 | 0.0015 |
| B-Monomer:Catalyst | 153.71 | 1 | 153.71 | 55.02 | < 0.0001 |
| C-Temperature | 0.0168 | 1 | 0.0168 | 0.006 | 0.9394 |
| D-Time | 0.0624 | 1 | 0.0624 | 0.0223 | 0.8835 |
| AB | 35.34 | 1 | 35.34 | 12.65 | 0.0035 |
| AC | 1.31 | 1 | 1.31 | 0.4693 | 0.5053 |
| AD | 2.04 | 1 | 2.04 | 0.732 | 0.4077 |
| BC | 0.3136 | 1 | 0.3136 | 0.1123 | 0.7429 |
| BD | 0.0182 | 1 | 0.0182 | 0.0065 | 0.9369 |
| CD | 12.01 | 1 | 12.01 | 4.3 | 0.0586 |
| $A^2$ | 0.1306 | 1 | 0.1306 | 0.0467 | 0.8322 |
| $B^2$ | 3.11 | 1 | 3.11 | 1.11 | 0.3108 |
| $C^2$ | 0.1306 | 1 | 0.1306 | 0.0467 | 0.8322 |
| $D^2$ | 0.0001 | 1 | 0.0001 | 0 | 0.995 |
| Residual | 36.32 | 13 | 2.79 | | |
| Lack of Fit | 31.77 | 10 | 3.18 | 2.09 | 0.2949 |
| Pure Error | 4.55 | 3 | 1.52 | | |
| Cor Total | 308.79 | 28 | | | |

**Table S23: PPS reduced model ANOVA of crude ester bond response**

| Source | Sum of Squares | df | Mean Square | F-value | p-value |
|---|---|---|---|---|---|
| Model | 234.15 | 3 | 78.05 | 31.24 | < 0.0001 |
| A-Anhydride:Epoxide | 45.09 | 1 | 45.09 | 18.05 | 0.0003 |
| B-Monomer:Catalyst | 153.71 | 1 | 153.71 | 61.51 | < 0.0001 |
| AB | 35.34 | 1 | 35.34 | 14.14 | 0.001 |
| Residual | 59.97 | 24 | 2.5 | | |
| Lack of Fit | 55.42 | 21 | 2.64 | 1.74 | 0.362 |
| Pure Error | 4.55 | 3 | 1.52 | | |
| Cor Total | 308.79 | 28 | | | |

**Table S24: PPS full model ANOVA of extract ester bond response**

| Source | Sum of Squares | df | Mean Square | F-value | p-value |
|---|---|---|---|---|---|
| Model | 218.38 | 14 | 15.6 | 4.69 | 0.0042 |
| A-Anhydride:Epoxide | 29.62 | 1 | 29.62 | 8.91 | 0.0105 |
| B-Monomer:Catalyst | 138.56 | 1 | 138.56 | 41.7 | < 0.0001 |
| C-Temperature | 0.066 | 1 | 0.066 | 0.0199 | 0.8901 |
| D-Time | 0.0296 | 1 | 0.0296 | 0.0089 | 0.9262 |
| AB | 21.07 | 1 | 21.07 | 6.34 | 0.0257 |
| AC | 2.89 | 1 | 2.89 | 0.8697 | 0.368 |
| AD | 1.72 | 1 | 1.72 | 0.5164 | 0.4851 |
| BC | 1.31 | 1 | 1.31 | 0.3945 | 0.5408 |
| BD | 0.0506 | 1 | 0.0506 | 0.0152 | 0.9037 |
| CD | 15.96 | 1 | 15.96 | 4.8 | 0.0472 |
| $A^2$ | 0.1237 | 1 | 0.1237 | 0.0372 | 0.85 |
| $B^2$ | 3.22 | 1 | 3.22 | 0.9685 | 0.343 |
| $C^2$ | 0.1541 | 1 | 0.1541 | 0.0464 | 0.8329 |
| $D^2$ | 0.1541 | 1 | 0.1541 | 0.0464 | 0.8329 |
| Residual | 43.2 | 13 | 3.32 | | |
| Lack of Fit | 39.41 | 10 | 3.94 | 3.12 | 0.1895 |
| Pure Error | 3.79 | 3 | 1.26 | | |
| Cor Total | 272.21 | 28 | | | |

**Table S25: PPS reduced model ANOVA of extract ester bond response**

| Source | Sum of Squares | df | Mean Square | F-value | p-value |
|---|---|---|---|---|---|
| Model | 205.3 | 6 | 34.22 | 12.77 | < 0.0001 |
| A-Anhydride:Epoxide | 29.62 | 1 | 29.62 | 11.05 | 0.0032 |
| B-Monomer:Catalyst | 138.56 | 1 | 138.56 | 51.7 | < 0.0001 |
| C-Temperature | 0.066 | 1 | 0.066 | 0.0246 | 0.8768 |
| D-Time | 0.0296 | 1 | 0.0296 | 0.011 | 0.9173 |
| AB | 21.07 | 1 | 21.07 | 7.86 | 0.0106 |
| CD | 15.96 | 1 | 15.96 | 5.95 | 0.0236 |
| Residual | 56.28 | 21 | 2.68 | | |
| Lack of Fit | 52.49 | 18 | 2.92 | 2.31 | 0.2681 |
| Pure Error | 3.79 | 3 | 1.26 | | |
| Cor Total | 272.21 | 28 | | | |

**Table S26: PCS full model ANOVA of extract $T_g$ response**

| Source | Sum of Squares | df | Mean Square | F-value | p-value |
|---|---|---|---|---|---|
| Model | 13117.63 | 14 | 936.97 | 15.34 | < 0.0001 |
| A-Anhydride:Epoxide | 0.6955 | 1 | 0.6955 | 0.0114 | 0.9168 |
| B-Monomer:Catalyst | 6183.53 | 1 | 6183.53 | 101.24 | < 0.0001 |
| C-Temperature | 341.36 | 1 | 341.36 | 5.59 | 0.0358 |
| D-Time | 2853.4 | 1 | 2853.4 | 46.72 | < 0.0001 |
| AB | 4.34 | 1 | 4.34 | 0.071 | 0.7944 |
| AC | 228.46 | 1 | 228.46 | 3.74 | 0.077 |
| AD | 0.0006 | 1 | 0.0006 | 9.63E-06 | 0.9976 |
| BC | 98.18 | 1 | 98.18 | 1.61 | 0.2289 |
| BD | 2975.38 | 1 | 2975.38 | 48.72 | < 0.0001 |
| CD | 41.77 | 1 | 41.77 | 0.684 | 0.4244 |
| $A^2$ | 3.38 | 1 | 3.38 | 0.0553 | 0.818 |
| $B^2$ | 34.57 | 1 | 34.57 | 0.566 | 0.4664 |
| $C^2$ | 3.13 | 1 | 3.13 | 0.0512 | 0.8248 |
| $D^2$ | 587.41 | 1 | 587.41 | 9.62 | 0.0092 |
| Residual | 732.92 | 12 | 61.08 | | |
| Lack of Fit | 697.23 | 9 | 77.47 | 6.51 | 0.0752 |
| Pure Error | 35.69 | 3 | 11.9 | | |
| Cor Total | 13850.55 | 26 | | | |

**Table S27: PCS reduced model ANOVA of extract $T_g$ response**

| Source | Sum of Squares | df | Mean Square | F-value | p-value |
|---|---|---|---|---|---|
| Model | 12627.45 | 5 | 2525.49 | 43.36 | < 0.0001 |
| B-Monomer:Catalyst | 6615.67 | 1 | 6615.67 | 113.59 | < 0.0001 |
| C-Temperature | 355.15 | 1 | 355.15 | 6.1 | 0.0222 |
| D-Time | 3040.54 | 1 | 3040.54 | 52.2 | < 0.0001 |
| BD | 3196.02 | 1 | 3196.02 | 54.87 | < 0.0001 |
| $D^2$ | 1919.92 | 1 | 1919.92 | 32.96 | < 0.0001 |
| Residual | 1223.1 | 21 | 58.24 | | |
| Lack of Fit | 1187.41 | 18 | 65.97 | 5.55 | 0.0917 |
| Pure Error | 35.69 | 3 | 11.9 | | |
| Cor Total | 13850.55 | 26 | | | |

**Table S28: PPS full model ANOVA of extract $T_g$ response**

| Source | Sum of Squares | df | Mean Square | F-value | p-value |
|---|---|---|---|---|---|
| Model | 293.12 | 14 | 20.94 | 6.02 | 0.0013 |
| A-Anhydride:Epoxide | 78.86 | 1 | 78.86 | 22.66 | 0.0004 |
| B-Monomer:Catalyst | 55.2 | 1 | 55.2 | 15.86 | 0.0016 |
| C-Temperature | 24.46 | 1 | 24.46 | 7.03 | 0.02 |
| D-Time | 0.9114 | 1 | 0.9114 | 0.2619 | 0.6174 |
| AB | 95.46 | 1 | 95.46 | 27.43 | 0.0002 |
| AC | 0.0227 | 1 | 0.0227 | 0.0065 | 0.9369 |
| AD | 8.61 | 1 | 8.61 | 2.47 | 0.1398 |
| BC | 14.8 | 1 | 14.8 | 4.25 | 0.0598 |
| BD | 0.1015 | 1 | 0.1015 | 0.0292 | 0.867 |
| CD | 7.41 | 1 | 7.41 | 2.13 | 0.1682 |
| $A^2$ | 0.0034 | 1 | 0.0034 | 0.001 | 0.9757 |
| $B^2$ | 4.92 | 1 | 4.92 | 1.41 | 0.2556 |
| $C^2$ | 0.1332 | 1 | 0.1332 | 0.0383 | 0.8479 |
| $D^2$ | 0.7745 | 1 | 0.7745 | 0.2226 | 0.6449 |
| Residual | 45.24 | 13 | 3.48 | | |
| Lack of Fit | 31.18 | 10 | 3.12 | 0.6653 | 0.727 |
| Pure Error | 14.06 | 3 | 4.69 | | |
| Cor Total | 339.11 | 28 | | | |

**Table S29: PPS reduced model ANOVA of extract $T_g$ response**

| Source | Sum of Squares | df | Mean Square | F-value | p-value |
|---|---|---|---|---|---|
| Model | 268.78 | 5 | 53.76 | 17 | < 0.0001 |
| A-Anhydride:Epoxide | 78.86 | 1 | 78.86 | 24.93 | < 0.0001 |
| B-Monomer:Catalyst | 55.2 | 1 | 55.2 | 17.45 | 0.0004 |
| C-Temperature | 24.46 | 1 | 24.46 | 7.73 | 0.0109 |
| AB | 95.46 | 1 | 95.46 | 30.18 | < 0.0001 |
| BC | 14.8 | 1 | 14.8 | 4.68 | 0.0417 |
| Residual | 69.58 | 22 | 3.16 | | |
| Lack of Fit | 55.52 | 19 | 2.92 | 0.6234 | 0.7784 |
| Pure Error | 14.06 | 3 | 4.69 | | |
| Cor Total | 339.11 | 28 | | | |
